# Flexible regions in the molecular architecture of Human fibrin clots structurally resolved by XL-MS and integrative structural modeling

**DOI:** 10.1101/739318

**Authors:** Oleg Klykov, Carmen van der Zwaan, Albert J.R. Heck, Alexander B. Meijer, Richard A. Scheltema

## Abstract

Upon activation, fibrinogen forms large fibrin biopolymers that coalesce into clots that assist in wound healing. Limited insights into their molecular architecture, due to the sheer size and insoluble character of fibrin clots, have however restricted our ability to develop novel treatments for clotting diseases. The so far resolved disparate structural details did provide insights into linear elongation; however, molecular details like the C-terminal domain of the *α*-chain, the heparin-binding domain on the *β*-chain, and others involved in lateral aggregation are lacking. To illuminate these dark areas, we applied crosslinking mass spectrometry (XL-MS) to obtain biochemical evidence in the form of over 300 distance constraints and combined this with structural modeling. These restraints additionally define the interaction network of the clots and e.g. provide molecular details for the interaction with Human Serum Albumin (HSA). We were able to construct the models of fibrinogen *α*(excluding two highly flexible regions) and *β*, confirm these models with known structural arrangements and map how the structure laterally aggregates to form intricate lattices together with fibrinogen *γ*. We validate the final model by mapping mutations leading to impaired clot formation. From a list of 22 mutations, we uncovered structural features for all, including a crucial role for *β* Arg’196 in lateral aggregation. The resulting model will be invaluable for research on dysfibrinogenemia and amyloidosis, as it provides insights into the molecular mechanisms of thrombosis and bleeding disorders related to fibrinogen variants. The structure is provided in the PDB-DEV repository.

---

Polymerization of fibrinogen is a multistage process that can be affected by numerous factors, including modifications of the initial building blocks, cellular environment, physiological properties of the reaction mixture and presence or absence of blood flow(1–3). Clot formation starts with the release of fibrinopeptides A and B from the N-termini of the *α*- and *β*-subunits of the large fibrinogen protein (Figure 1A). Fibrinopeptide release is followed by the A:a knob-hole interaction of the parallel fibrinogens and simultaneous linear elongation through the sewing of the *γ*-nodules with the next fibrinogen hexamer(4–6) (Figure 1B). Less is known about the next step, consisting of lateral aggregation of fibrin oligomers or protofibrils (Figure 1C). Protofibrils have successfully been visualized with microscopy methods(7, 8) and their packing into fibrin fibers with a periodicity of 225 Å shown(9, 10), although this so far provided only a low-resolution view. This final, 3-dimensional scaffold is formed through bi-lateral or tri-molecular junction mechanisms(11, 12) and some potential sites involved in lateral aggregation are shown in Figure 1E. Concurrently with the polymerization process, covalent crosslinking of fibrin mediated by Factor XIIIa occurs(13), which strengthens the final structure with naturally occurring disulfide bonds(14). For a detailed review about clot formation see Weisel & Litvinov(15). The clots are finally degraded by plasmin cleavage upon formation of a ternary complex combining the clot, t-PA and plasminogen (Figure 1C,D) followed by conversion of plasminogen into plasmin(16). As much as currently is understood about these processes, the molecular structure of the full fibrinogen molecules and how they form clots is currently lacking. To date, it has been difficult to capture the structure with traditional structural biology techniques like Crystallography or EM and structural models are solely available for smaller disconnected parts. To illustrate, a high-resolution crystal structure was reported(17); however, this lacks domains important for lateral aggregation like the *α*C-terminal, *β* N-terminal and *γ*C-terminal domains.

Crosslinking mass spectrometry (XL-MS) has emerged as a powerful tool for studying protein-protein interactions. It is capable of extracting protein structure distance information from any protein regardless of size, solubility or strength of interaction(18) and has found application to tissue samples(19). The technique utilizes small reagents with two reactive moieties that form a covalent bond between two amino acids in close proximity. Upon application to and proteolytic digestion of protein-protein complexes four distinct peptide products have formed: non-modified, mono-linked, loop-linked and crosslinked peptides. The first three classes consist of single peptides in various forms that yield no or limited structural information. The fourth group consists of two peptides bound together by the crosslinking reagent; this yields valuable distance information for protein tertiary structure (the two peptides originate from the same protein) or for protein quaternary structure (the two peptides originate from different proteins). Identification of the two peptides by mass spectrometry allows for localization of the crosslink within the proteins at the residue level and uncovers distance constraints useful for assembling a model of the protein complex under investigation. We apply XL-MS utilizing DSSO (a lysine-lysine reactive crosslinking reagent) to study fibrin clots extracted from platelet poor plasma and uncover 284 unique distance constraints. This dataset was combined with structural modeling (protein structure prediction and protein-protein docking; combined with XL-MS this approach is termed integrative structural modeling) to illuminate dark areas on the fibrinogen structure. We now, supported by biochemical evidence, resolve 78 % (previously 66 %, counted by amino acids present in the structure) of the total fibrin structure after processing, validating 92 % of the distance constraints in our dataset. With the illuminated regions, lateral aggregation within fibrin clots can be visualized at the molecular level. To validate the final structure, we investigated 22 mutations reported to lead to impaired clot formation. For the complete set, our model provides explanations at the molecular level. Additionally, it was in other studies suggested that albumin interferes with fibrinolysis by interfering with activation or hindering access to cleavage sites of clot degradation enzymes(20). From our data, we can decipher the interaction interface between albumin and fibrin clots. This shows that albumin interferes with fibrinolysis by sterically hindering access to sites involved in plasminogen activation and plasmin cleavage.

**Figure 1.**
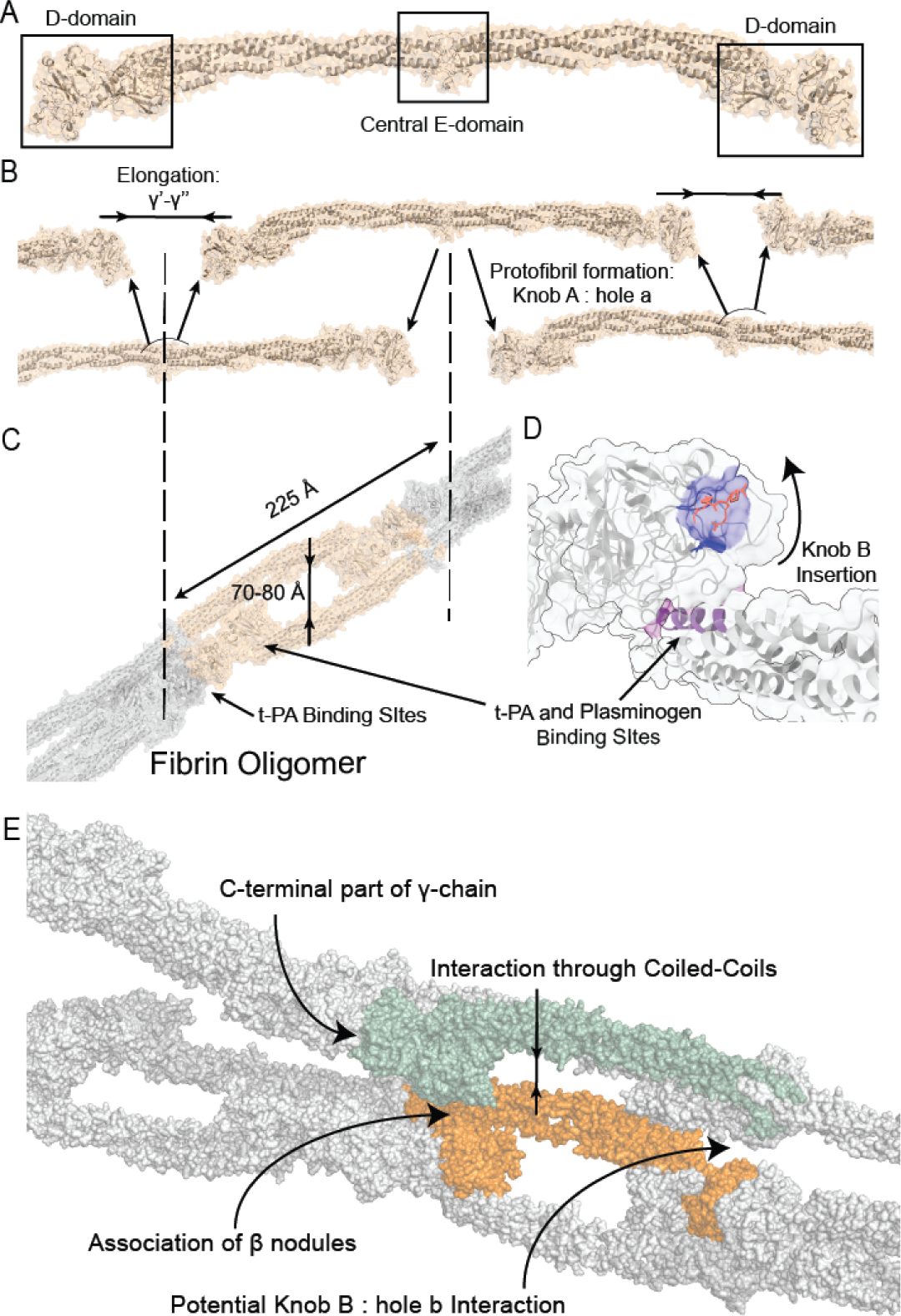
Structural model of fibrin clot formation. **(A)** Structural model of the fibrinogen hexamer building block (based on PDB: 3ghg). **(B)** The hexamer building blocks form protofibrils through linear elongation via the well-described A:a knob-hole interaction. **(C)** Formation of the fibrin oligomer. **(D)** The potential function of knob B upon insertion into hole B. **(E)** Lateral aggregation of the protofibrils, including interaction through laterally aggregating coiled-coils, association of *β*-nodules and the putative site of the knob b: hole B interaction.

## Materials and Methods

For fibrinogen, we adopt the residue numbering scheme as provided in the Uniprot database, i.e. based on the numbering in the nascent form of the individual chains including the signal peptides and fibrinopeptides.

### Fibrin clot sample preparation

Plasma samples were obtained after informed consent and in accordance with the ethics board of Sanquin (Amsterdam, The Netherlands). Three samples of blood extracted from one voluntary donor, citrated to prevent coagulation, were treated separately to induce clot formation and consecutively to crosslink the structure. After a series of centrifugation steps at 120, 2000, and 10000 g clot formation in 1 mL of platelet poor plasma was initiated by addition of 16 mM CaCl2 (Sigma-Aldrich) and 9 pM tissue factor innovin (Siemens Healthcare Diagnostics). The final mixtures were heated to 37 °C for 5 min, the resulting clots were washed with crosslinking buffer containing 20 mM HEPES, 150 mM NaCl, and 1.5 mM MgCl2 (all Sigma-Aldrich) adjusted to pH 7.8 with NaOH (Sigma-Aldrich) and with no reagent present. This buffer is designed to preserve the native conformation of the proteins, and therefore the clot, while washing removes unspecifically bound proteins. Crosslinking buffer was aspirated from the clots, which were consequently filtered with a molecular weight cut-off (MWCO) filter from Amicon (10 kDa, Sigma-Aldrich), Microcon (30 kDa, Merck Millipore) and one sample was processed without prior filtering. After these steps, crosslinking buffer supplemented with 2 mM DSSO(18) (Thermo Fisher Scientific) was pushed through the filters in four repetitions to maintain an ample supply of fresh non-degraded crosslinking reagent. The crosslinking reaction was then quenched by addition of a 20 mM Tris-HCl (Sigma-Aldrich) solution at pH 8.5 and the crosslinked clots were then snap frozen in liquid nitrogen.

The snap-frozen crosslinked clots were processed as previously described with minor adjustments(18). Briefly, clots were homogenized on a bead-mill (Retsch) for 5 min at 120 Hz. Then the samples were treated with protein deglycosylation mix II (NEBB) overnight, attempting to fully remove both N-as well as O-linked glycans, followed by acetone-cold precipitation to clean up the protein mixture. As deglycosylation is applied after the crosslinking reaction, the removal of the glycans will not impact the distance information derived from the crosslinks while improving the identification performance. Proteins were resuspended in a solution containing 1 % SDC and 10 mM TCEP with 40 mM CAA (Sigma-Aldrich) as reduction and alkylating agents and heated to 37 °C for 1 hour. The resulting solution was diluted with 50 mM ammonium bicarbonate (Sigma-Aldrich) and digested by a combination of LysC (Wako) and Trypsin (Promega). The final peptide mixtures were desalted with BioSelect solid-phase extraction C18 columns (Vydac) and fractionated with an Agilent 1200 HPLC pump system (Agilent) coupled to an Opti-LYNX trap column (Optimize Technologies) and SCX-separation column (PolyLC), resulting in 25 fractions per fibrin clot. For the serum experiments, human serum from an anonymous healthy donor was provided by Sanquin Research (Amsterdam, The Netherlands). The whole blood was collected in a 9-mL Vacuette tube (Greiner Bio-One) containing Z Serum Clot Activator and then was left undistributed at r.t. for 30-60 min. The clotted material was removed by centrifugation at 1800 g for 20 min at r.t and the sera was transferred as a 1 mL aliquot to a clean 1.5 mL Eppendorf tube, snap frozen in liquid nitrogen and stored at −80 °C until further analysis. Proteins were processed in a similar fashion as the clots, excluding homogenization, deglycosylation and fractionation.

### Liquid chromatography with mass spectrometry and data analysis

Each fraction was separated with an Agilent 1290 Infinity uHPLC system (Agilent) on a 50 cm analytical column packed with C_18_ beads with 300 Å pore-size coupled online to an Orbitrap Fusion Lumos Tribrid mass spectrometer (Thermo Fisher Scientific). For the serum experiments, the Orbitrap HF-X (Thermo Fisher Scientific) was used with the same acquisition parameters. Details of the full setup, separation gradient, data acquisition methods, and data analysis were described previously(18). For the shotgun proteomics experiments the same approach was used except that unfractionated samples were used for analysis. Quantification was performed with the iBAQ (intensity-Based Absolute Quantification) algorithm in MaxQuant(21). For analysis of the specificity of HSA binding, quantification of detected precursor ions was performed across all the available crosslinked fractions. From the 1205 unique crosslink pair identifications, we retained only those detected in two out of the three samples, resulting in 451 unique peptide residue pairs (from 697 spectral matches), 284 of which (from 437 spectral matches) provide distance constraints for the fibrinogen subunits and their interacting proteins. As each sample is treated slightly different prior to crosslinking, the selected crosslinks represent the core interaction points. Furthermore, as the sample is solid-phase which decreases the space between atoms when compared to solution, combined with the mechanical forces applied during clot purification, we define the maximum crosslinker distance as 43 Å. This is in line with previously published studies applying solid-phase crosslinking(19) and crosslink-driven docking(22). The RAW data and all associated output files used in this study have been deposited to the Proteome-Xchange Consortium via the PRIDE partner repository with the identifier PXD011680.

### Modeling and docking of albumin and individual fibrinogen domains into fibrin

Predictions of the location of individual domains were performed with the web-services provided by ThreaDomEx(23) and Robetta(24). Additional template searches were performed with HHalign-Kbest(25). Each domain was modeled separately with I-TASSER(26), Robetta(24) and RaptorX(27). A highly detailed description of these modeling procedures, allowing for replication of the results, is provided in Supplementary Note 1. The resulting structures were scored with z-DOPE(28), Errat(29), Procheck(30), QMEAN(31), ProQ2(32) and the final model of each domain chosen based on a combination of available scores (best or second best in at least 3 out of 5 scores), combined with previously reported biochemical features of the modeled domain to ensure biological validity. Tables with the scores are provided in Supplementary Note 1. The detected intralinks within each domain were used for validation of the resulting model (apart from *α*432-491, where the crosslinks were not used in the validation procedure).

Interlinks and overlength intralinks (potentially arising from the interaction between two copies of the same subunit) were clustered with DisVis(33) to determine collections of crosslinks pointing to each location. Clusters generated with DisVis are provided as Supplementary Figures 4-8. Disvis input parameters are provided in Supplementary Files 1. For docking laterally aggregating protofibrils from the template (PDB: 3ghg), we split the fibrinogen hexamer to a single trimer. Trimers were submitted to CPORT(34) to highlight active residues, which were included in the docking procedure. Docking was performed with HADDOCK(35) and ClusPro(36). Highly detailed descriptions of each docking run, allowing for replication of the results, are provided in Supplementary Notes 2 and 3. Docking parameters including defined restraints can be found in Supplementary Files 2 and 3. Obtained structures were visualized with pyMol(37) and ChimeraX(38). The assembled model of fibrin clots is deposited into the PDB-Dev database with identifier PDB-DEV 00000030. Docking results of clots to albumin are provided as Supplementary Models 1-3.

### Analysis of mutation sites

Conservation analysis of the generated structures has been performed with the Consurf webserver(39) with standard settings using 300 sequences from the Uniref-90 database(40) and modeled structures as PDB input. Mapping of the conservation sites and relevant tables can be found in Supplementary Data 1-8. CADD(41) scores were calculated with the web version of ANNOVAR(42) with the reported mutations as input (for further details see http://wannovar.wglab.org/; higher scores are more likely to have detrimental effects). A CADD score represents an aggregation of multiple scoring methods operating at the nucleotide level, describing the effect of a genetic mutation on the protein of interest and therefore serves as an effective parameter to highlight deleterious, functional, and/or disease causative variants. For those residues with reported mutations, the side chain backbone-dependent rotamers were generated with pyMol and the conformation with the lowest strain values and no clashes within the structure selected for final analysis.

## Results

So far studies aimed at deciphering the high-resolution structure of fibrin assemblies used a combination of atomic force microscopy and molecular dynamics simulations(43, 44). These studies provide invaluable insights into the structural organization of fibrin oligomers, but to circumvent technological limitations were not performed on full clots. A recent study mapped endogenously crosslinked peptide species detected by mass spectrometry, which suggested a more compact organization of protein blocks then so far assumed(45) and, importantly, highlights the potential of mass-spectrometry based investigations of fibrin clots. Here we employ *in situ* crosslinking mass spectrometry on purified blood clots to specifically delve into the mechanisms underlying lateral aggregation (Supplementary Figure 2).

**Figure 2.**
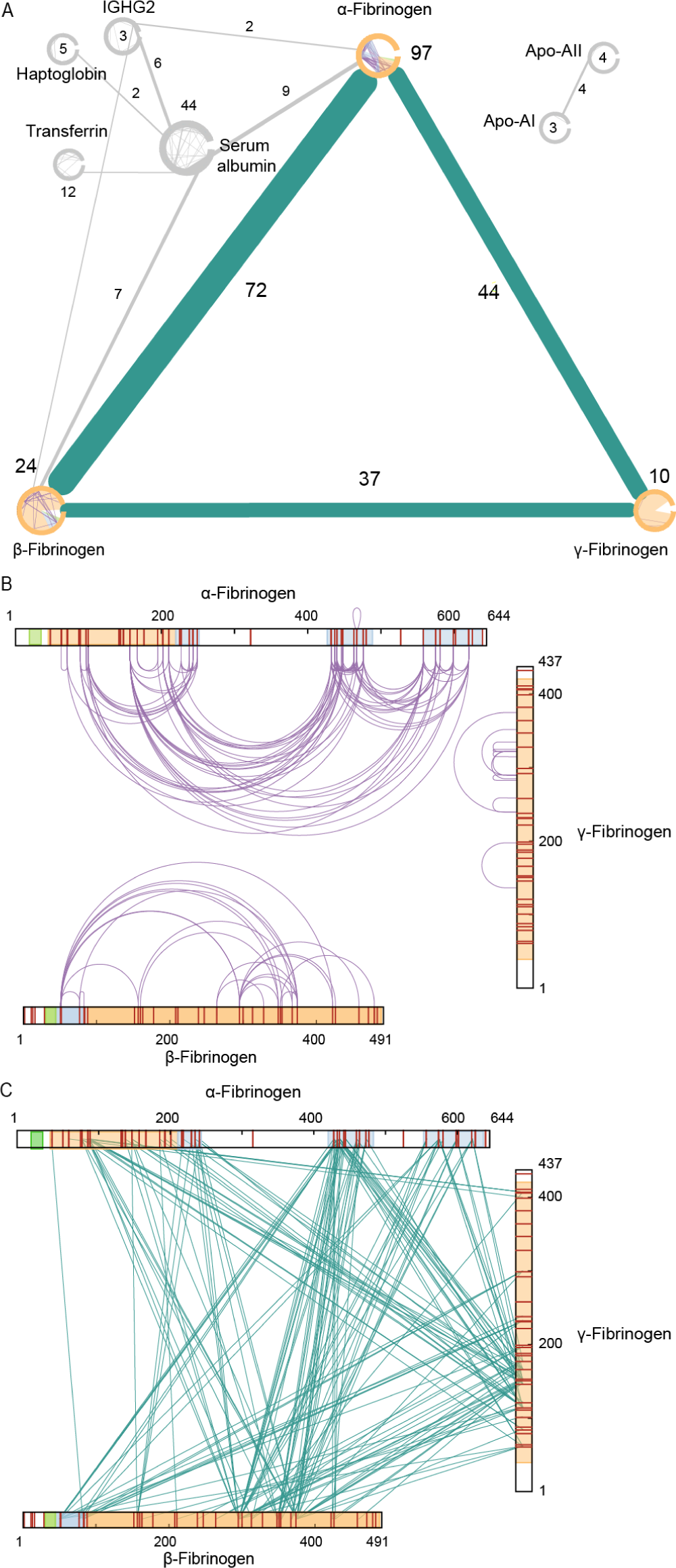
Overview of crosslinking MS data. **(A)** Overview of crosslinks detected in 2 out of 3 samples. Only proteins with at least 1 inter- and 1 intralink are depicted. **(B)** XL-MS intra- and **(C)** interlinks detected on the sequences of the fibrinogen chains. Cleaved off fibrinopeptides A and B are shown in green and domains with reported PDB structures are shown in orange. Domains for which additional structural models are presented in this study are shown in blue.

### Initial data check

To gain insight into the sample purity and data quality of our XL-MS measurements, we first visualized the detected crosslinks as a network(46). We only retained proteins with at least 1 intra- and 1 interlink (Figure 2A) to filter out for which insufficient structural information is detected. The core of the resulting network is built up from the fibrinogen *α*-(97 intralinks; 116 interlinks to the other chains), *β* - (24 intralinks; 109 interlinks to the other chains), and *γ*-chain (10 intralinks; 81 interlinks to the other chains). In addition to crosslinks between the fibrinogen subunits, we also detect crosslinks involving Serum albumin, Transferrin, Haptoglobin, IgGs, and Apolipoproteins. In total three interlinks were detected between fibrinogen and Immunoglobulin, while for HSA 44 intralinks and 16 interlinks to the *α*- and *β*-chains of fibrinogen were detected. The locations of the intra- and interlinks show the data covers a very large degree of the sequence of fibrinogen subunits (Figure 2B and C). As anticipated, for two lysine poor regions on the *α*-chain no crosslinks were detected.

### Mapping detected restraints on available structures

For structural validation, we mapped the detected intralinks of Serum albumin on the existing crystal structure, for which we find 90.9 % are within the set distance constraint (PDB: 1uor; Supplementary Table 1). For fibrinogen, the detected inter- and intralinks were mapped on existing crystal structures of the fibrinogen hexamer and fibrin(4, 17). For the first partial structure of fibrin (PDB: 1fzc) a total of 43 could be mapped of which 81 % were within the allowed distance constraint. For the assembled *α, β*, and *γ* trimer (PDB: 3ghg), used as the basis for this study, a total of 86 crosslinks could be mapped of which 58 % were within the allowed distance constraint (Supplementary Table 2). The lower percentages within distance crosslinks indicate that upon assembly of the full protofibril likely some degree of higher order organization or polymerization takes place. These details were so far not captured as the previous studies focused on individual regions of the full structure.

**Table 1.** Overview of fibrinogen domains.

| Fibrinogen subunit | Residue numbers | Comment | Available structure | Submitted to structure prediction | Involved in detected crosslinks |
| --- | --- | --- | --- | --- | --- |
| $\alpha$ | 1-19 | Signal Peptide | No | No | No |
|  | 20-35 | Fibrinopeptide A | No | No | No |
|  | 36-45 | No structural info | No | No | No |
| | 46-249 | α-chain coiled-coil | Partially, 46-219 (PDB: 3ghg) | Yes, 220-249 | Yes |
|  | 250-431 | Includes highly flexible parts | No | No | No |
|  | 432-491 | Part of N-term subdomain (Interactive Domain) | Full template (PDB: 2jor, based on the structure of bovine domain) | Yes | Yes |
|  | 492-557 | Includes highly flexible parts | No | No | No |
|  | 558-620 | Part of C-term subdomain (RGD-Containing Domain) | No | Yes | Yes |
|  | 621-629 | No structural info | No | No | No |
|  | 630-644 | Cleaved during processing | No | No | No |
| $\beta$ | 1-30 | Signal Peptide | No | No | No |
|  | 31-44 | Fibrinopeptide B | No | No | No |
|  | 45-51 | No structural info | No | No | No |
|  | 51-83 | Part of knob B (Heparin-Binding Domain) | No | Yes | Yes |
| | 88-486 | β-chain coiled-coil and knob | Yes (PDB: 3ghg) | No | Yes |
| $\gamma$ | 1-26 | Signal Peptide | No | No | No |
|  | 27-38 | No structural info | No | No | No |
| | 39-420 | γ-chain coiled-coil and nodule | Yes (PDB: 3ghg) | No | Yes |
|  | 421-450 | No structural info | No | No | No |

**Table 2.** Analyzed fibrinogen mutation sites.

| Gene | Amino acid change | Associated to / source | CADD score | Conservation score | Potential effect | Conservation scores of affected residues |
| --- | --- | --- | --- | --- | --- | --- |
| FGA | P571H | Amyloidosis(55) | 17.0 | 4 | Change of hydrophobic Pro'571 to polar Histidine - affects flexibility through e.g. π-π stacking with Phe'570 or cation-π interaction with Arg'573. The Histidine gives rise to a H-π interaction with Ser'576 and interferes with an important structural function implied by its high conservation score. Alternatively, a H-π interaction with Glu'240 is also possible. Another rotamer provides the potential for π-π stacking with Tyr'601 fixing an otherwise flexible loop into beta-sheet extension. | αPhe'570-1<br>αSer'576-9<br>αTyr'601-1 |
| FGA | R573C,<br>R573H,<br>R573L | Dysfibrinogenemia(58, 59),<br>Amyloidosis(60) | 18.2,<br>13.4,<br>13.7 | 6 | In αC1, αArg'573 is involved in a H-H bond with the side chain of αLys'225 in the folded conformation. In other organisms, the Arg is often changed to Ser or Thr, also capable of maintaining this interaction. Within αC2, this mutation has the potential to form a salt bridge with αGlu'447. Mutation to Histidine provides the potential for π-π stacking to the neighboring αPhe'570 and potentially a cation-π interaction to αLys'227. Amyloidosis-causing substitution to Leucine provides the potential for α580-603 beta-sheet extension. | αLys'225-9,<br>αGlu'447-6,<br>αPhe'570-1,<br>αLys'227-1,<br>αArg'573-6 |
| FGB | R196C | Dysfibrinogenemia(61) | 28.9 | 6 | In addition to the formation of albumin-complexes, βArg'196 is crucial for lateral aggregation through a salt bridge with βGlu'177. In some organisms, this residue is changed to Lys, which also allows for salt bridge formation. | βGlu'177-8 |

### Illuminating dark areas on the fibrinogen structure

The available structures for fibrinogen and fibrin lack several domains, or as we term them dark areas. To define those dark areas amenable to integrative structural modeling supported by our XL-MS results, we separated the chains into domains for which currently no structure is available (Table 1). In total, we identified four suitable domains. (1) For the N-terminal domain of the *β*-chain at residue 51-83, 29 inter- and two intralinks were detected. This domain was previously reported to bind the anticoagulant Heparin(47) and even though a heparin binding pocket is not strictly defined, some structural characteristics are present(48, 49). A total of 18 models were chosen for manual inspection. By enforcing the exposure of a heparin-binding site and the restraints imposed by the intralinks, a high scoring conformation could be selected (Figure 3A; Supplementary Table 2, Beta within). (2) The *α*-chain domain at residue 220-249 was detected with 4 intra- and 19 interlinks. From a total of 15 models, we selected two likely models with different conformations (folded and elongated) that score high on all metrics and satisfy the detected intralinks well (in both cases 4/4 valid restraints; Supplementary Table 2, Alpha1). As the two versions are likely present (see next paragraph), we selected both (Figure 3B). (3) For the *α*-chain domain at residue 432-491 (part of the *α*C-terminal domain), we detected 69 inter- and 20 intralinks. This domain was previously reported to be involved in fibrin clot formation by interacting with other fibrin copies, making it unclear for the detected intralinks whether they arise from intra- or inter-molecular contacts. A structure for the bovine form was previously reported (PDB: 2jor)(50), which we used for homology modeling with no distance restraints but with forced disulfide bond formation between *α*Cys’461 and *α*Cys’491. It was possible to map all 20 detected restraints within the set distance constraint for the best scoring of 15 models (Figure 3C; Supplementary Table 2, Alpha3a Within). (4) For the *α*-chain domain at residue 558-620, we detected 50 inter- and seven intralinks. Within this domain, upon activation three lysines are hypothesized to interact with glutamine residues, either between different molecules or with the flexible region of the *α*C-terminal domain, forming *β*-sheets(51). From 16 manually inspected models, we chose the highest scoring conformation with an exposed integrin-binding RGD motif fixed in place by the *β*-sheets (Figure 3D; Supplementary Table 2, Alpha3b within).

**Figure 3.**
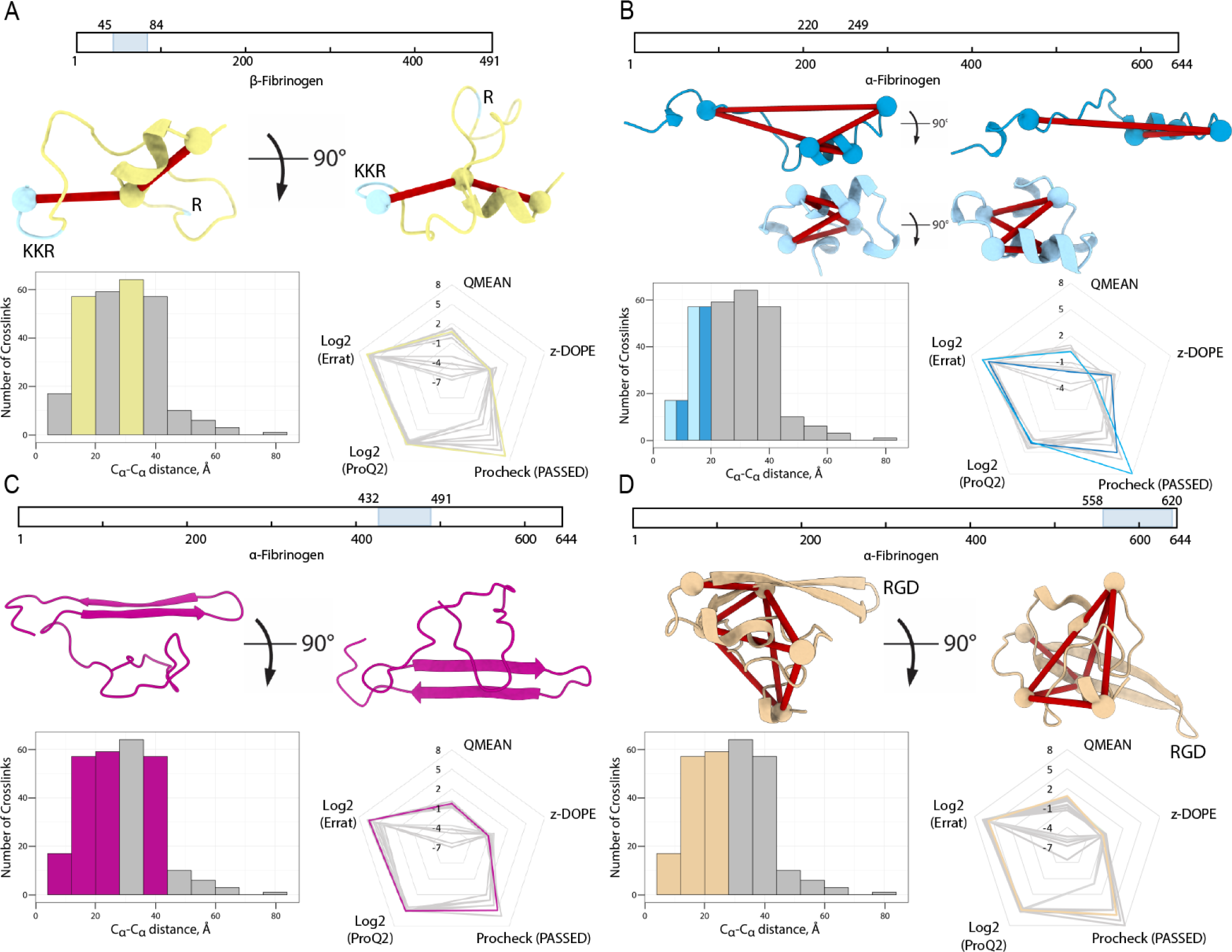
Structural models of individual domains. **(A)** The *β*-fibrinogen N-terminus, residues 51-83. **(B)** Part of the *α*-fibrinogen domain 1, residues 220-249. **(C)** The N-terminal subdomain of the *α*-fibrinogen C-terminus, residues 432-491. **(D)** The C-terminal subdomain of the *α*-fibrinogen C-terminus, residues 558-620.

**Figure 4.**
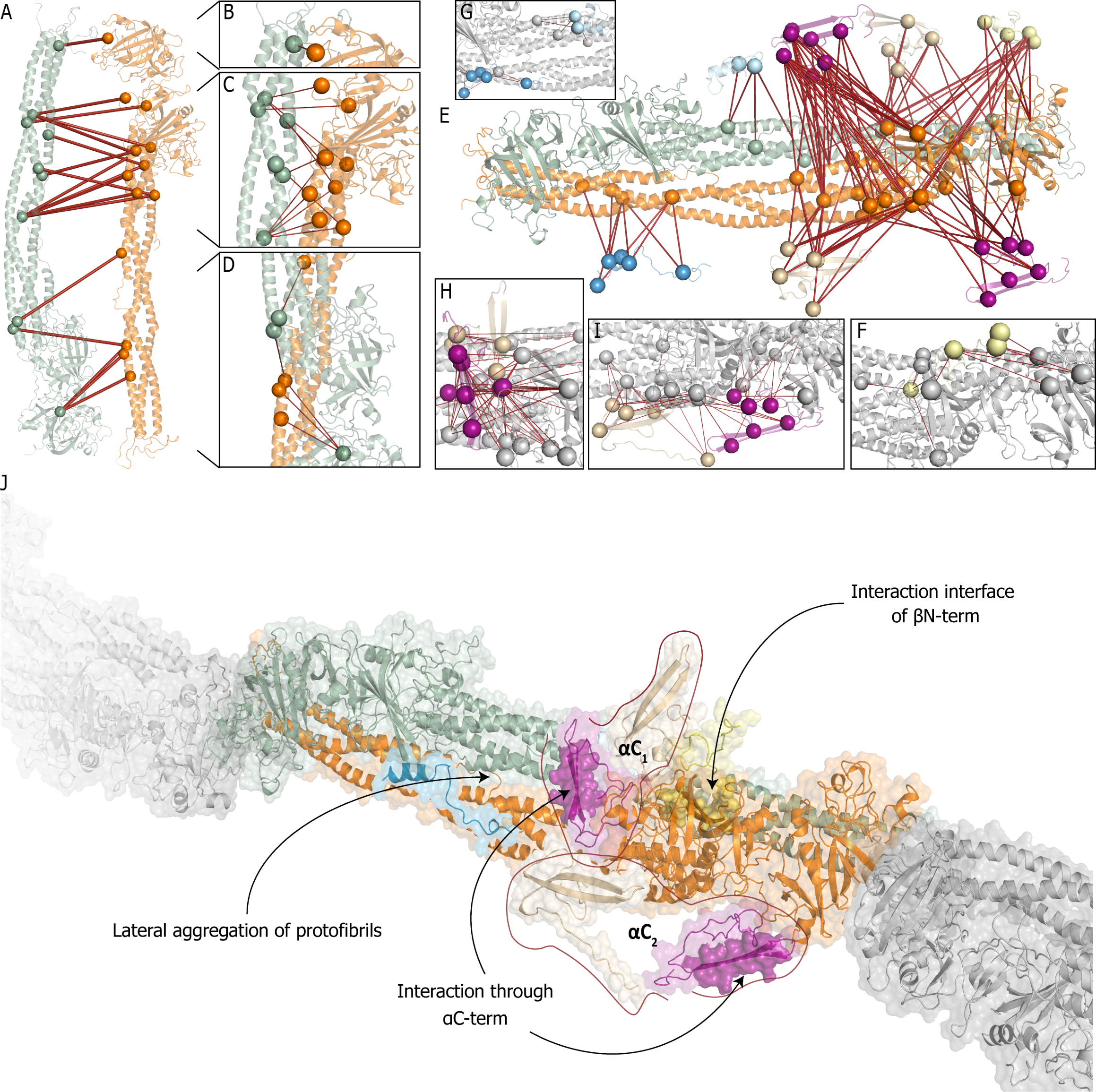
Assembly of the full fibrin clot model. **(A)** Overview of detected crosslinks in the fibrin scaffold with one protofibril shown in green and another in orange. **(B-D)** Zoom-in to crosslink-rich regions on the assembled structural model with the zoom on top **(B)**, middle **(C)** and bottom **(D)** parts of the docked scaffold. **(E)** Placement of the structural models of previously not-resolved domains which are generated in this work. **(F-H)** Zoom-in to crosslink-rich regions on the assembled model with the scaffold (grey): **(F)** *β* N-termini (yellow), **(G)** *α*-fibrinogen residues 220-249 in folded (light blue) and elongated (blue) conformations, (H) *α*-Interactive domain Cluster I (purple, top) and RGD-binding domain Cluster II (beige, top), and **(I)** *α*-Interactive domain Cluster II (purple, bottom) and RGD-binding domain Cluster I (beige, bottom). **(J)** The final model of fibrin with the highlighted interaction interfaces with the color scheme described above. Each copy of *α*-interactive and -RGD domains are grouped as they belong to the same fibrinogen molecule.

The domains at residues 1-35 on the *α*-chain, 1-44 on the *β*-chain and 1-26 on *γ*-chain were excluded as they harbor signal- and/or fibrinopeptides, which were also not detected in our shotgun proteomics experiments (Supplementary Figure 3). Residues 36-45 on the *α*-chain, 45-50 and 85-87 on the *β*-chain, 27-38 and 421-437 on the *γ*-chain were omitted, as we detected no structural information for these regions. The domain 250-431 on the *α*-chain (previously reported to be highly flexible(9)) was excluded as we do not detect structural information, while the remaining part before *α*250 represents the naturally occurring fibrinogen truncated variants (i.e. Fragment X)(52). The domain at residue 492-557 on the *α*-chain (part of the *α*C-terminal domain, but preceding the domain we uncovered in 4) was excluded as we also did not detect structural information; this domain was reported to potentially form beta-sheets upon activation by FXIII(51), which we cannot confirm from our data as there are no crosslinks detected. These preceding regions are considered as highly flexible linkers in our model. The *α*-fibrinogen C-terminal domain at residues 621-644 was excluded as we also did not detect structural information for this domain; in line with previous reports that it is partly removed post-translationally(53). In our shotgun proteomics experiment, we detect peptides from the elongated 420-kDa version of *α*-chain; however, these peptides comprise 0.3 % of the total detected intensity and we expect them not to play an important role (Supplementary Figure 3). In total, we improve the fibrinogen structure coverage, supported by biochemical evidence, from 66 % to 78 %.

### Assembly of the fibrin scaffold

The assembly of the final repeating unit was driven by a combination of our XL-MS results and interaction interface analysis by DisVis and CPORT for scaffold assembly and finally protein-protein docking with HADDOCK (Figure 4). As the initial fibrinogen template possess C2 symmetry, we split the hexamer into pairs of identical trimers. CPORT analysis shows that most of the potential interaction sites are localized on discrete parts of the molecule and are not shared between the trimers. From our DisVis analysis, we identified two major clusters of overlength intralinks between the fibrinogen chains (Cluster I: 24; Cluster II: 7 and 5 unclustered crosslinks; Supplementary Figure 4). With the set of crosslinks from Cluster I, protein docking with HADDOCK on the two copies of the fully assembled fibrinogen trimer was performed to create the protofibril that acts as the basis for the next steps (Figure 4A). After docking, 18/24 crosslinks were within the defined distance constraint (Figure 4B-D; note that HADDOCK utilizes the distance constraints only as a post-docking filter; Supplementary Table 2, dimer). Interestingly, 5 out of 6 overlength crosslinks and all crosslinks from Cluster II together with 5 unclustered restraints involve residues *β* Lys’295 and *β* Lys’374 on the *β*-nodule, which together with the predicted interaction interfaces from DisVis suggests they arise from the antiparallel alignment of the *β*-nodules

To place the modeled domains onto the final structure, we perfomed HADDOCK docking with the interlinks from each domain as the final filtering step (Figure 4E). (1) From 29 detected interlinks for the *β* N-terminus, eight crosslinks are used later for placement of the *α*C-terminus domains and the remaining 21 are used for placement of the *β* N-terminus onto the dimer. All 21 crosslinks are identified as true positives by the DisVis analysis (Supplementary Figure 5). After docking, 16/21 crosslinks were within the set distance constraint (Figure 4F; Supplementary Table 2, Beta withDimer); the overlength crosslinks are however located on a flexible region in the *β*-subunit and can be explained from this perspective. (2) For the *α*-chain domain at residue 220-249, 23 interlinks were detected to the other chains. Both the elongated as well as the folded conformation of this domain were fused to *α*His’219, with additional refinement to optimize placement accuracy and side-chain placement (Figure 4G). After docking, 19/23 interlinks for the elongated conformation and 4/23 for the folded conformation were within the set distance constraint (Supplementary Table 2, Alpha1a Elongated and Alpha1a Folded). As the folded conformation hides accessible lysines, the imbalance of explained interlinks was anticipated. (3) The C-terminus of the *α*-chain was previously suggested to be split into an N- and C-terminal subdomain(51); this is in line with our domain predictions (Table 1). For the *α*-chain domain at residue 432-461 (N-terminal sub-domain), 69 interlinks were detected to the *α*-chain; the large number of interlinks was expected due to its highly interactive nature(51). From our DisVis analysis, we detected three clusters of interlinks (Cluster I: 46, Cluster II: 19, and Cluster III: 4; Supplementary Figure 6), indicating this domain is localized in three distinct positions in our sample. Cluster I is placed in front of the *β* nodule (Figure 4H, purple residues). After docking 44/46 inter-links are within the set distance constraint (Supplementary Table 2, Alpha3a 01). Cluster II is placed under the *β* and *γ* nodules (Figure 4I, purple residues). After docking 19/19 interlinks are within the set distance constraint (Supplementary Table 2, Alpha3a 02). Cluster III is placed in the middle of the coiled-coil region of the aggregated protofibrils (Supplementary Table 2, Alpha3a Central), which we hypothesize arises from a transitional state towards the active form(9). We note, however, that with four crosslinks this is an observation of relatively low confidence and we do not pursue this further. (4) For the *α*-chain domain at residue 558-620 (C-terminal subdomain), 50 interlinks were detected of which 17 are to the indistinguishable N-terminal sub-domains and therefore not included in the docking procedure. The remaining crosslinks are to the other chains and therefore provide localization information for the C-terminal subdomain. Fitting with 3, from our DisVis analysis we also detect two clusters of interlinks (Cluster I: 21, Cluster II: 12; Supplementary Figure 7), indicating that both this domain as well as the N-terminal sub-domain are localized in two distinct positions in our samples. After docking, 18/21 interlinks for Cluster I (Figure 4I, beige; Supplementary Table 2, Alpha3b 01) and 7/12 interlinks for Cluster II were within the set distance constraint (Figure 4H, beige; Supplementary Table 2, Alpha3b 02). Supporting this placement, the 17 discarded crosslinks are within the set distance constraints when mapped between the N- and C-terminal sub-domains after docking (Supplementary Table 2, Alpha3a 3b). Based on this observation, we suggest that each C-terminal sub-domain can be matched to its respective N-terminal sub-domain, the combinations of which we term *α*C1 and *α*C2 (see Figure 4J). Outliers from each docking run were mapped on the alternative subunit (where applicable), resulting in two additional valid restraints.

**Figure 5.**
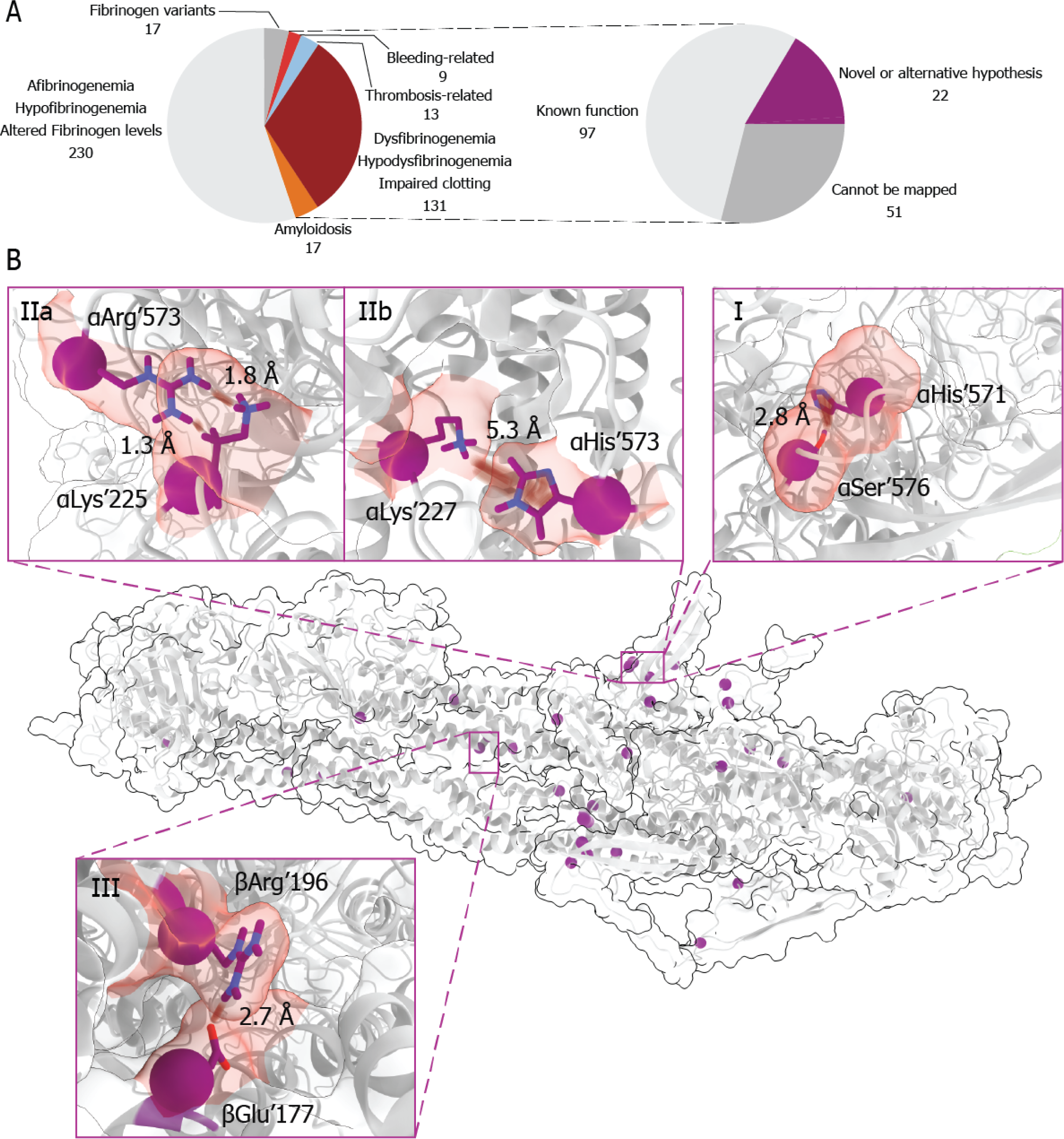
Mapping known mutations on the final structural model of fibrin clots. **(A)** Selection of mutations of interest from previously reported mutations. **(B)** Locations of selected mutation sites (purple). Panels I-IV show several highlighted mutations. Panel I showcases the *α*His’571 in interaction with *α*Ser’576. Panel IIa shows the role of *α*Arg’573 and Panel IIb shows an additional interaction with *α*Lys’227 created by its substitution to *α*His’573. Panel III depicts salt bridge formation between *β* Arg’196 and *β* Glu’177 from two laterally aggregated protofibrils, which is disrupted when *β* Arg’196 is mutated to Cys.

**Figure 6.**
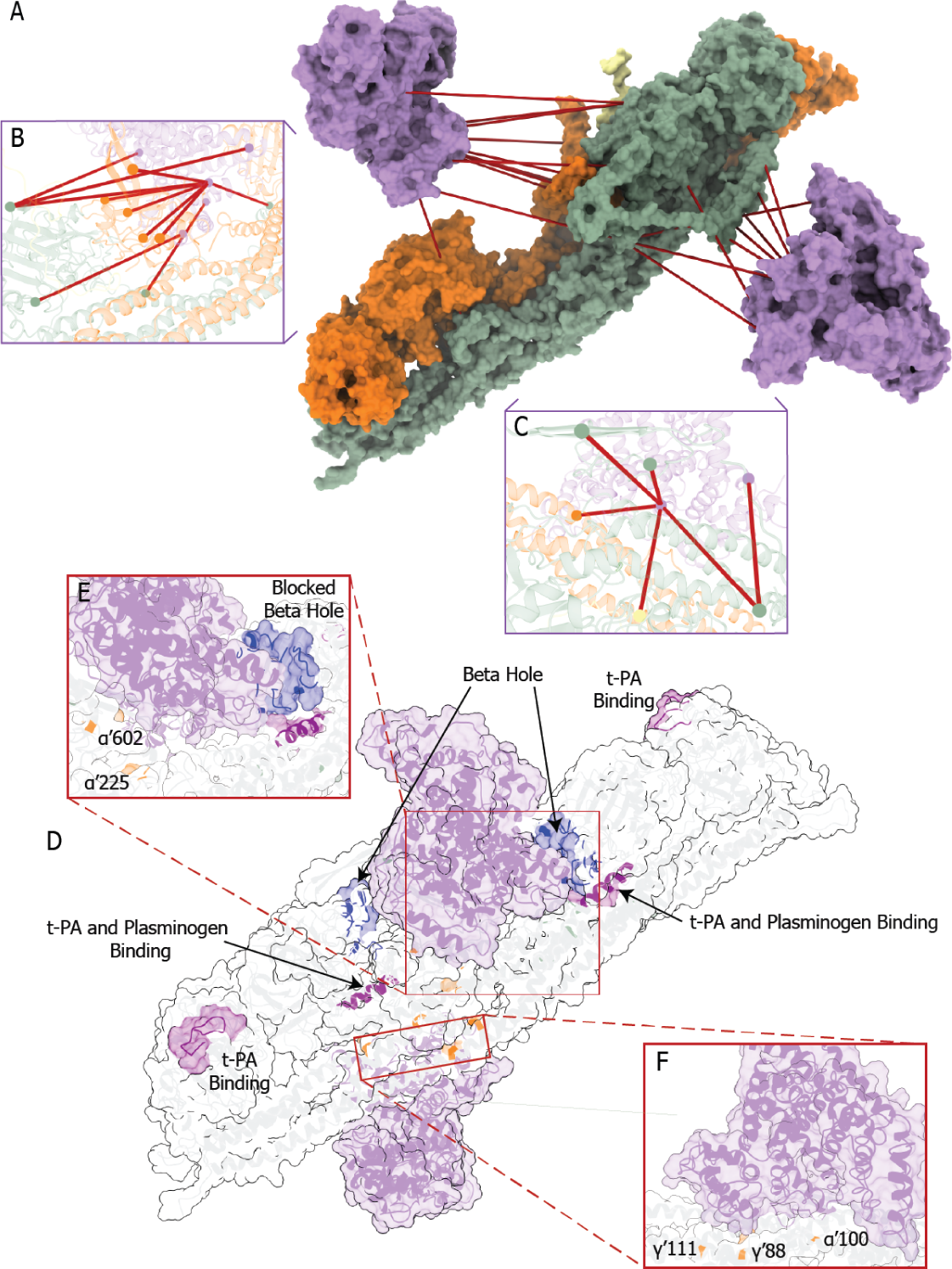
Interaction Interfaces between the fibrin clot and HSA. **(A)** Overview of the detected crosslinks between albumin and the fibrin clot. **(B, C)** Placement of albumin according to Cluster I and Cluster II restraints on the fibrin clot. Only validated restraints are shown. **(D)** Delaying fibrin degradation by hindering access to sites of ternary complex formation and plasmin cleavage with a zoom-in of **(E)** Cluster I and **(F)** Cluster II.

### Interaction Interface of albumin and fibrin clots

As the fibrin trimer is present in 2 copies, a total of 31 crosslinked pairs (of 16 detected) can be mapped, for which we used a structure of HSA without ligand (PDB: 1uor). DisVis analysis uncovered 3 distinct clusters of 17, 9 and 4 crosslinks respectively (Supplementary Figure 8). One not clustered restraint, as well as restraints from Cluster III, represents the mirrored location of Clusters I and II respectively and were not used in docking. The selected docked model, obtained with restraints from Cluster I, places albumin inside the cavity on the structural model of the fibrin clot (Figure 6A, left). It is possible to map 11 out of 17 restraints within the defined distance cut off (Figure 6B, Supplementary Model 1). The selected model, obtained with restraints from Cluster II, places albumin on the exposed part of the fibrin clot (Figure 6A, right) with 6 valid restraints out of 9 (Figure 6C, Supplementary Model 2). In conclusion, from 16 detected crosslinks it was possible to map 13 as valid (Supplementary Table 3). Interestingly, 3 outliers can be placed on another model docked from Cluster I (Supplementary Model 3). This model possesses higher balanced scores but in total satisfies only 9 restraints and therefore was not selected (Supplementary Note 3).

## Discussion

Fibrin clot formation has received a large amount of interest over the years and at a molecular level many details have been resolved through various structural biology techniques. From the generated partial protein structures, for example, details on how fibrinogen elongates were readily uncovered(4–6, 54). Fibrinogen however also laterally aggregates to form thick fibers and intricate lattices, the molecular details of which are less well understood. We applied a combination of XL-MS and structural modeling to these dark regions and shed light on the mechanisms underlying lateral aggregation. This approach is not hampered by protein complexity and/or size and flexibility, albeit it provides structural information of low resolution. From the presented dataset of 284 distance constraints, it was not only possible to validate structural predictions but also unambiguously place them on the full structure, for which the final model validates 92 % of the detected distance constraints. Because XL-MS is not limited to the proteins of interest, we also uncover interactors (like HSA and IgG). Based on sufficient amounts of detected inter protein crosslinks, we were also able to dock HSA on our final fibrin model and uncovered a surprising role for HSA. As the used docking software uses the distance constraints only as a final filtering step and we ensured that the structural prediction software did not include this data where possible, the distance constraints provided by XL-MS constitute an independent biochemical validation for this structure.

### Mapping known mutations

The assembled structural model is only useful if it provides insights into the molecular details explaining how mutations result in impaired clot formation (Figure 5). From a list of 417 known mutations, we focused on 174 mutations potentially related to altered clot structure, including 22 mutations for which our new structural model provides insights (Figure 5a). From the final list of 170, 57 % could already be explained and 30 % were not possible to map on our structure (four had no reported sites; 24 are located on the *α* chain, before residue 46, and 2 on the *β* chain, before residue 51, both of which are part of signal peptides and fibrinopeptides; 16 are located on flexible domains, and finally five arise from frameshift mutations on the *β* and *γ* C-terminal domains). Additionally, we examined an overlapping prediction list of 304 mutations with calculated CADD scores(41), and combine it with our calculated conservation scores(39) (Supplementary Data 1-8). For all of the selected 18 mutation sites, it was possible to uncover molecular details explaining impaired clot formation (Supplementary Table 4). Here we discuss in more detail a subset of three mutations (Figure 5b; Table 2). (I) For *α*Pro’571 mutated to Histidine (CADD score: 17.0), we find the potential for a hydrogen-*π* interaction with the highly conserved *α*Ser’576 (Figure 5b; panel I). Previously, this mutation was linked to amyloidosis(55), hypothesized to occur through beta-sheet extension in the region *α*580-603(56, 57). Interaction to *α*Ser’576 observed in our structure confirms that the Histidine introduces additional stiffness in the domain causing extensive beta-sheet formation in the region. The high conservation score of *α*Ser’576 additionally indicates an important role for this residue in normal clots, which with the hydrogen-*π* interaction will be reduced or even abolished. (II) For *α*Arg’573 three debilitating mutations were so far reported(58–60). Even though this residue is only moderately conserved (score 6 out of 9), from our structure we find that *α*Arg’573 from the RGD-containing domain in *α*C1 is involved in an H-H bond to the highly conserved residue *α*Lys’225 in the folded conformation of the *α*220-249 domain. The moderate conservation score mostly arises from the replacement of Arginine to Serine or Threonine in other organisms, which are also capable of forming an H-H bond to *α*Lys’225. This bond makes it structurally important for the correct folding of *α*558-620 and its loss alone potentially disturbs correct clot formation (Figure 5b; panel IIa). Mutation to Cysteine (CADD score: 18.2; fibrinogen Dusart) is associated with impaired clot formation by the involvement of the Cysteine with albumin complexes through S-S bonds(58). Cysteine is a polar residue also capable of forming H-H bonds and this substitution does not necessarily affect the structure directly. Mutation to Histidine (CADD score: 13.4; fibrinogen San Diego V), also associated with impaired clot formation, has so far not satisfactorily been explained. From our model, we find the potential of *α*His’573 to form a cation-*π* interaction with *α*Lys’227 (Figure 5b; panel IIb). This Lysine is involved in the correct folding of the *α*220-249 domain and might disturb the dynamic equilibrium between the elongated and folded forms. Loss of this equilibrium affects the correct placement of these subunits that are directly involved in lateral aggregation to *α*C1 and *α*C2. This fits well with previous reports that clots with decreased turbidity values and relatively fewer laterally aggregated protofibrils are found for fibrinogen San-Diego V(59). The role of the mutation to Leucine (CADD score: 13.7) remains elusive, but we hypothesize that this affects the flexibility of the region in *α*558-620 resulting in beta-sheet extension. (III) For *β* Arg’196 mutated to Cysteine (CADD score: 28.9; fibrinogen Longmont), we find in our model that the Arginine is crucial for lateral aggregation of protofibrils through the coiled-coil region by the formation of a salt bridge to *β* Glu’177 on the parallel coil (Figure 5b; panel III). Its loss disrupts this process, supported by the finding that polymerization is not normalized after removal of fibrinogon albumin complexes(61).

### Interaction of fibrin clots with Serum albumin

There are several proteins which were previously reported as non-covalently bound to fibrin clots(20, 62, 63) and almost all of them are presented in our shotgun proteomics data (Supplementary Data 9). After filtering our crosslinking dataset we only retain restraints for albumin, Apolipoproteins, and Immunoglobulins (Supplementary Data 10). Although we detected Apolipoproteins in our dataset, there is no intermolecular connection to fibrin. In the case of Immunoglobulins, the amount of detected restraints is relatively low and all are focused on a single immunoglobulin residue, making prediction of the interaction interface extremely difficult. In the case of albumin, 16 intermolecular restraints were detected allowing placement of the albumin on fibrin clots with high confidence (Figure 6A-C). Before proceeding, we evaluated the specificity of this binding by estimating albumin content in both human serum as well as in our clot samples (Supplementary Figure 9A,B). Generally, albumin is several folds more abundant than fibrinogen in plasma (64). This situation alters drastically in our clot samples, where only a 1.75-fold increase of albumin is observed (Supplementary Figure 2B). We also examined the presence of albumin in human serum and it is still the most abundant protein with several folds excess over most detected proteins. Further confirmation of the specificity of the interaction was derived from the detected abundances of albumin intra- and inter-molecular crosslinks to the fibrin clot (Supplementary Figure 9C). As it is generally harder to form an interlink, it is expected that interlinks have lower detected abundances. In our data, intra- and interlinks however have the same abundance, further confirming the specificity of the interaction at the indicated stoichiometry.

Fibrin clots are cleaved by plasmin, which is formed upon activation of plasminogen. Activation takes place after formation of a ternary complex comprising of t-PA, plasminogen, and fibrin. The first cleavage points of plasmin are located at 3 sites on the C-terminal domain(65). Then so-called “D-dimers” are formed, which consists of two connected D-domains and 1 E-domain(16, 66). Products of fibrin degradation can be further cleaved into smaller pieces with the cleavage on multiple sites(65, 67). Two sites on fibrinogen are known to bind either t-PA or both t-PA and plasminogen. Previously, it was suggested that upon clot formation a structural rearrangement takes place that makes both sites active(68). More specifically, site *α*’167-179 is normally buried inside the structure and exposed after insertion of knob B into hole b(69). In our model, the placement of albumin suggested by Cluster I blocks hole B (Figure 6D,E). This placement interferes with knob B insertion and therefore blocks the site for ternary complex formation. It was also reported that *α*C-termini contain cryptic t-PA and plasminogen high-affinity binding sites that are active only in fibrin clots. However, the exact location of these sites remains elusive but are likely between *α*’444 and *α*’619. In our structural model, access to a large part of *α*C1 is hindered by albumin. We hypothesize that placement within this cavity explains the decrease of fibrinolysis rates elevated albumin concentrations. In addition, apart from interfering with ternary complex formation, albumin hinders access to a number of plasmin cleavage sites (Figure 6D, Supplementary Table 5). The location suggested by Cluster I ensures that albumin covers site *α*’602 from *α*C1 (Figure 6E), which has been reported to be cleaved by plasmin(65). The location suggested by Cluster II ensures that albumin binding interferes with cleavage by blocking plasmin cleavage sites *α*Arg’123 and *γ*Lys’88 completely(67) (Figure 6F). Another cleavage site is located on *γ*Lys’111, however, cleavage here is known to be less efficient which is in line with our structural model where access to that site is hindered by the laterally aggregating regions within the fibrin clot. In total, we docked two albumin molecules to the structural model of a fibrin clot. If we take into account all 31 possible crosslinked peptide pairs between the fibrin clot and HSA, we could also mirror the model from Cluster II. Such a placement results in binding of three albumin molecules to six copies of each fibrinogen protein (Supplementary Figure 10). This is indeed very close to our copy-number estimation (Supplementary Figure 2B), which suggests a ratio of 1.76 to 1 of albumin to fibrinogens.

### Binding of fibrin clots to endothelial cells and thrombocytes

Excitingly, our structural model also provides a detailed molecular basis for the binding sites for platelets and endothelial cells. The definition of the *α*-chain domain at residue 558-620 with an RGD domain fixed in place by beta-sheets, biochemically supported by 50 inter- and seven intralinks both in terms of location within the clot structure as well as the folding of the domain. This provides an excellent starting point for further structural investigations. Several important binding events are associated with precisely this spot due to the affinity to Integrin receptors exposed on the cellular membrane(70–72). In the case of platelet binding, this RGD-motif competes with *γ*-fibrinogen and the interaction between fibrin and integrin *α*IIb*β* 3 was previously proposed as a therapeutic target to destabilize thrombi(73, 74). We envision that XL-MS can be used to study the interactions made by both platelets as well as endothelial cells to provide a molecular picture for these interactions. Potentially, from such an unbiased screen more information in terms of other receptors and/or other locations on the full fibrin clot structure can be obtained. Importantly though, from the detailed structural information, this will open up avenues for the development approaches for the treatment of thrombosis and/or hemophilia.

## Acknowledgments

We thank all members of the Heck-group for their helpful contributions, especially Kelly Dingess for support with the video. We acknowledge financial support by the large-scale proteomics facility Proteins@Work (Project 184.032.201) embedded in the Netherlands Proteomics Centre and supported by the Netherlands Organization for Scientific Research (NWO). Additional support came through the European Union Horizon 2020 program FET-OPEN project MSmed (Project 686547) and the European Union Horizon 2020 program INFRAIA project Epic-XS (Project 823839).

## Contributions

R.A.S. conceived of the study. R.A.S., A.B.M., and A.J.R.H. designed the experiments. C.v.d.Z. performed clotting and clot clean up; O.K. performed clot processing, data analysis, and structural investigations; O.K. and R.A.S. wrote the manuscript. All authors critically reviewed and edited the manuscript.

