## Supplementary Material for "Flexible regions in the molecular architecture of Human fibrin clots structurally resolved by XL-MS and integrative structural modeling"

**Running title:** Intra- and inter-molecular architecture of fibrin clots

Oleg Klykov^a,b^, Carmen van der Zwaan^c^, Albert J.R. Heck^a,b^, Alexander B. Meijer^a,c^, Richard A. Scheltema^a,b,1^

^a^ Biomolecular Mass Spectrometry and Proteomics, Bijvoet Center for Biomolecular Research and Utrecht Institute for Pharmaceutical Sciences, Utrecht University, Padualaan 8, 3584 CH Utrecht, The Netherlands.

^b^ Netherlands Proteomics Centre, Padualaan 8, 3584 CH Utrecht, The Netherlands.

^c^ Department of Molecular and Cellular Hemostasis, Sanquin Research, Amsterdam 1066 CX, The Netherlands

Richard A. Scheltema, Utrecht University, Padualaan 8, 3584 CH Utrecht, The Netherlands. tel: +31 30 253 6793

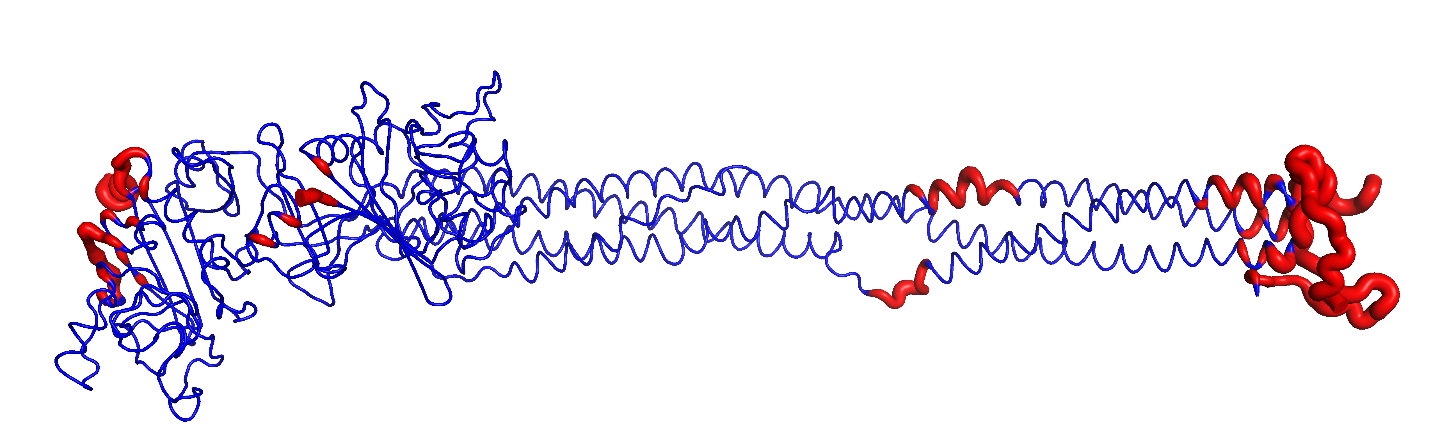

**Supplementary Figure 1:** CPORT prediction of interaction interfaces of the fibrinogen trimer to other copies of the trimer (PDB: 3ghg; marked in red). Highlighted regions on the outer edges were previously reported to be involved in linear elongation. The highlighted region within the β-knob was previously hypothesized to be involved in protofibril formation, while the highlighted region on the coiled-coil is potentially involved in lateral aggregation.

**Supplementary Table 1:** Mapping of detected crosslinks on a known HSA structure (PDB: 1uor). Only restraints detected in two out of three samples are retained.

| **SequenceA** | **PositionA** | **SequenceB** | **PositionB** | **Score** | **# Csms** | **Type** | **Distance, Å** |
| --- | --- | --- | --- | --- | --- | --- | --- |
| RHPYFYAPELLFFA[K]R | 183 | Y[K]AAFTECCQAADK | 186 | 206.51 | 4 | Intra | 4.61 |
| L[K]CASLQK | 223 | ASSA[K]QR | 219 | 97.91 | 2 | Intra | 7.55 |
| LVNEVTEFA[K]TCVADESAENCDK | 75 | F[K]DLGEENFK | 36 | 142.53 | 3 | Intra | 9.45 |
| LVTDLT[K]VHTECCHGDLLECADDR | 264 | AF[K]AWAVAR | 236 | 127.32 | 2 | Intra | 10.69 |
| LVRPEVDVMCTAFHDNEETFL[K]K | 160 | Y[K]AAFTECCQAADK | 186 | 78.1 | 2 | Intra | 10.69 |
| AEFAEVS[K]LVTDLTK | 257 | AF[K]AWAVAR | 236 | 74.83 | 5 | Intra | 10.70 |
| QNCELFEQLGEY[K]FQNALLVR | 426 | [K]QTALVELVK | 549 | 122.5 | 5 | Intra | 11.04 |
| CC[K]HPEAK | 463 | NLG[K]VGSK | 456 | 67.38 | 1 | Intra | 11.30 |
| VGS[K]CCK | 460 | HPEA[K]R | 468 | 55.06 | 1 | Intra | 11.50 |
| NYAEA[K]DVFLGMFLYEYAR | 347 | LA[K]TYETTLEK | 375 | 91.27 | 6 | Intra | 12.64 |
| [K]VPQVSTPTLVEVSR | 438 | AT[K]EQLK | 565 | 52.59 | 2 | Intra | 12.96 |
| LDELRDEG[K]ASSAK | 214 | NLG[K]VGSK | 456 | 85.18 | 2 | Intra | 13.34 |
| RHPYFYAPELLFFA[K]R | 183 | ECCE[K]PLLEK | 305 | 100.27 | 7 | Intra | 13.79 |
| LA[K]TYETTLEK | 375 | AF[K]AWAVAR | 236 | 134.12 | 1 | Intra | 13.88 |
| NLG[K]VGSK | 456 | ASSA[K]QR | 219 | 44.95 | 1 | Intra | 14.09 |
| VT[K]CCTESLVNR | 499 | CASLQ[K]FGER | 229 | 72.82 | 1 | Intra | 14.71 |
| VGS[K]CCK | 460 | ASSA[K]QR | 219 | 47.04 | 2 | Intra | 14.83 |
| LDELRDEG[K]ASSAK | 214 | VGS[K]CCK | 460 | 74.26 | 1 | Intra | 15.34 |
| ADLA[K]YICENQDSISSK | 286 | F[K]DLGEENFK | 36 | 69.43 | 1 | Intra | 16.33 |
| EFNAETFTFHADICTLSE[K]ER | 543 | NLG[K]VGSK | 456 | 82.14 | 1 | Intra | 16.64 |
| EQL[K]AVMDDFAAFVEK | 569 | [K]QTALVELVK | 549 | 80.56 | 3 | Intra | 17.54 |
| AAFTECCQAAD[K]AACLLPK | 198 | [K]YLYEIAR | 161 | 105.82 | 2 | Intra | 17.61 |
| EFNAETFTFHADICTLSE[K]ER | 543 | LDELRDEG[K]ASSAK | 214 | 157.34 | 2 | Intra | 19.22 |
| VT[K]CCTESLVNR | 499 | LA[K]TYETTLEK | 375 | 98.13 | 2 | Intra | 19.31 |
| LA[K]TYETTLEK | 375 | L[K]CASLQK | 223 | 74.85 | 1 | Intra | 19.49 |
| QNCELFEQLGEY[K]FQNALLVR | 426 | AT[K]EQLK | 565 | 54.59 | 4 | Intra | 19.70 |
| VT[K]CCTESLVNR | 499 | L[K]CASLQK | 223 | 74.42 | 1 | Intra | 20.02 |
| [K]QTALVELVK | 549 | CC[K]ADDK | 584 | 73.97 | 1 | Intra | 20.94 |
| SLHTLFGD[K]LCTVATLR | 97 | F[K]DLGEENFK | 36 | 77.75 | 1 | Intra | 21.32 |
| L[K]CASLQK | 223 | VGS[K]CCK | 460 | 71.48 | 2 | Intra | 21.46 |
| SLHTLFGD[K]LCTVATLR | 97 | [K]YLYEIAR | 161 | 89.78 | 2 | Intra | 23.30 |
| VFDEF[K]PLVEEPQNLIK | 402 | AF[K]AWAVAR | 236 | 74.6 | 1 | Intra | 24.34 |
| VFDEF[K]PLVEEPQNLIK | 402 | VT[K]CCTESLVNR | 499 | 110.35 | 1 | Intra | 24.97 |
| F[K]DLGEENFK | 36 | [K]YLYEIAR | 161 | 89.78 | 3 | Intra | 25.10 |
| VT[K]CCTESLVNR | 499 | AT[K]EQLK | 565 | 59.89 | 1 | Intra | 26.43 |
| VT[K]CCTESLVNR | 499 | [K]YLYEIAR | 161 | 120.78 | 1 | Intra | 36.44 |
| AAFTECCQAAD[K]AACLLPK | 198 | F[K]DLGEENFK | 36 | 60.96 | 2 | Intra | 37.34 |
| AVMDDFAAFVE[K]CCK | 581 | [K]YLYEIAR | 161 | 69.44 | 1 | Intra | 39.33 |
| VT[K]CCTESLVNR | 499 | ETCFAEEG[K]K | 597 | 64.55 | 1 | Intra | 39.96 |
| ETCFAEEG[K]K | 597 | [K]YLYEIAR | 161 | 60.96 | 2 | Intra | 45.88 |
| VT[K]CCTESLVNR | 499 | F[K]DLGEENFK | 36 | 95.9 | 1 | Intra | 46.79 |
| TYETTLE[K]CCAAADPHECYAK | 383 | [K]YLYEIAR | 161 | 82.14 | 1 | Intra | 53.04 |
| VFDEF[K]PLVEEPQNLIK | 402 | F[K]DLGEENFK | 36 | 74.34 | 1 | Intra | 53.15 |

**Supplementary Table 2:** Full list of detected crosslinks with distances mapped on the fibrin clot structure. Columns correlate to chains and their respective residues, mapped distance in Å, full protein names and includes comments on which part of the fibrin clot structure mapping is performed.

See the excel sheet ‘Supplementary Table 2.xlsx’.

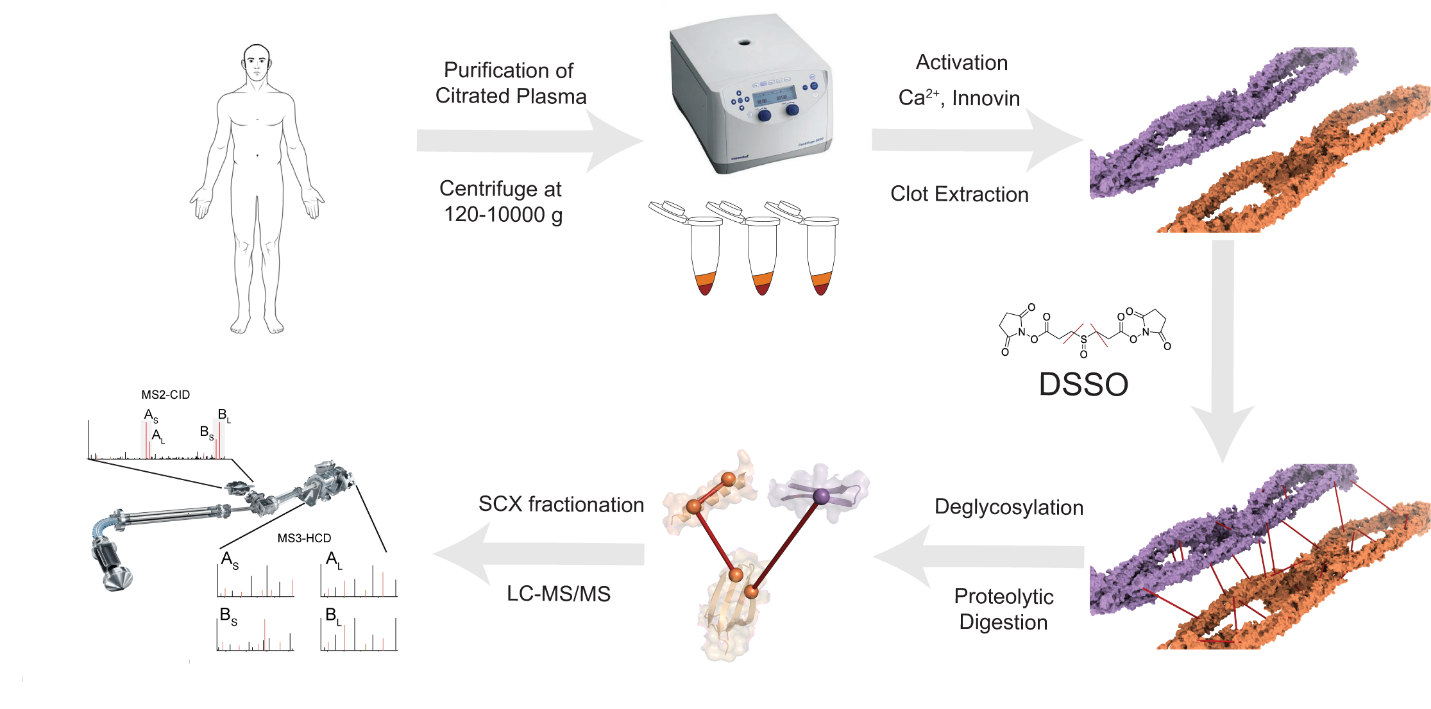

**Supplementary Figure 2:** Workflow for preparation and processing of crosslinked fibrin clots. **(A)** Plasma samples were centrifuged in three steps at 120-10000 g to obtain platelet-poor plasma. **(B)** Clot formation was induced with Ca^2+^ and innovin. Clots were extracted with MWCO filters and **(C)** crosslinked with DSSO. **(D)** Samples were processed according to our sample preparation protocol with additional deglycosylation. **(E)** Resulting peptide mixtures were fractionated with SCX and then subjected to LC-MS/MS analysis.

**Supplementary Note 1:** Modelling of individual subunits of fibrinogen.

**βN-term (Fibrinogen β’51-84)**

*Submitted Amino Acid Sequence:*

KKREEAPSLRPAPPPISGGGYRARPAKAAATQKK

Initially, modelling of the whole β-fibrinogen was attempted, because a template (PDB: 3ghg) is available that covers most of the sequence (**ROBETTA Comparative as part of whole Fibrinogen Beta**). The resulting model however did not conform to our quality criteria and we continued with modelling of region 51 – 84 for which crosslinks were detected. ITASSER modeling with no template was performed in parallel with comparative Robetta (**ITASSER and ROBETTA Comparative single piece**). Additionally, ITASSER outputs were used as templates in crosslink-driven refinement (**Template-based ITASSER**) and the best models were examined. The Raptor X model was excluded as it did not satisfy one of the 2 detected crosslinks within this domain (**Raptor X**). The remaining models were in line with the detected restraints and were analyzed with several available scoring algorithms. Importantly, preservation of residues from the exposed basic pocket reported to bind heparin(1) was observed in all models. As a result, model 2 from **Robetta Comparative (single piece)** was selected as the final model.

R

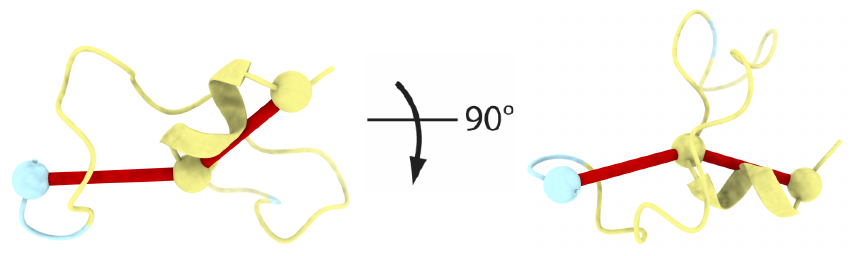

R

KKR

KKR

*Crosslinked residues:*

| **Residue A** | **Residue B** |
| --- | --- |
| 77 | 51 |
| 77 | 83 |

*Generated models:*

| **Model number** | **Satisfied Restraints**  **(out of 2)** | **ProQ2** | **z-DOPE Score** | **ProCheck**  **(Error/Warning/Pass)** | **QMEAN** | **Errat**  **Overall Quality Factor** |
| --- | --- | --- | --- | --- | --- | --- |
| **ITASSER with no template** |  |  |  |  |  |  |
| 1 | 2 | 20.16 | -0.75 | 5/4/0 | -4.21 | 77 |
| 2 | 2 | 26.98 | -0.48 | 5/3/1 | -6.72 | 66 |
| 3 | 2 | 22.38 | 0.24 | 6/2/0 | -6.17 | 91 |
| 4 | 2 | 25.96 | -0.65 | 6/2/0 | -4.95 | 97 |
| 5 | 2 | 16.88 | -0.28 | 6/3/0 | -6.72 | 74 |
| **Robetta Comparative**  **(as part of whole Fibrinogen Beta)** |  |  |  |  |  |  |
| 1 | 2 | 13.37 | -0.46 | 1/2/5 | 0.16 | 82 |
| 2 | 2 | 24.88 | -0.52 | 0/2/7 | 0.95 | 73 |
| 3 | 2 | 19.33 | -0.32 | 0/2/7 | 0.36 | 77 |
| 4 | 2 | 17.14 | -0.00 | 1/4/4 | 0.07 | 64 |
| 5 | 2 | 20.43 | -0.38 | 0/2/6 | 1.05 | 94 |
| **Raptor X** |  |  |  |  |  |  |
| 1 | 1 | n/a | 0.01 | n/a | n/a | n/a |
| **Robetta Comparative**  **(single piece)** |  |  |  |  |  |  |
| 1 | 2 | 22.85 | -0.64 | 2/1/5 | 0.39 | 86 |
| **2** | **2** | **27.56** | **-0.90** | **0/1/7** | **0.74** | **100** |
| 3 | 2 | 22.54 | -1.06 | 2/2/4 | 1.29 | 74 |
| 4 | 2 | 23.00 | -0.92 | 0/1/7 | 1.08 | 94 |
| 5 | 2 | 17.10 | -0.86 | 0/2/6 | -0.22 | 94 |
| **Template-based ITASSER** |  |  |  |  |  |  |
| ITASSER_Model 1 as template | 2 | 27.35 | -0.43 | 3/3/3 | -3.02 | 52 |
| ITASSER_Model 2 as template | 2 | 31.90 | 0.42 | 5/1/2 | -4.53 | 8 |

**The α-fibrinogen domains.**

Before modelling, we split this protein on individual domains based on detected crosslinks, ThreaDomEx software and domain predictions from the Robetta server.

| **ThreaDomEx** | **Robetta** | **Available Structure** | **Submitted to structure prediction after examining XL positions** |
| --- | --- | --- | --- |
| 45-172; 173-224; | 46-227; 228-292 | Partially, 3ghg | 220-249 |
| 225-425 | 293-415 | No | No |
| 426-555; 556-644 | 416-510; 511-630 | Partially, not experimental | As 2 separate domains, 431-491 and 558-620 |

**1^st^ α-chain Domain (α-fibrinogen’220-249)**

*Submitted Amino Acid Sequence:*

HLPLIKMKPVPDLVPGNFKSQLQKVPPEWK

Our model for this domain is an extension of an existing template (PDB: 3ghg). First ITASSER, *ab initio* Robetta, Raptor (**ITASSER, Robetta msa, Raptor X**) and additional ITASSER refinement with template-based and restraint-driven modelling were performed (**Template-based ITASSER**). All of the models satisfied the defined distance cut off for the restraints. Modelling of this part was performed after finalization of the scaffold (Supplementary Note 2). Therefore, there was an opportunity to estimate how well crosslinks to this scaffold fit when the model is fused to the scaffold. For all the models, two border conformations were automatically generated: Folded and Elongated. For the Elongated conformation, it was possible to define the best model based on additional scoring (Robetta msa Model 5). For the Elongated conformation, the best model was selected based on the mapped restraints to the scaffold. Both models after refinement with ITASSER satisfy 22 out of 23 restraints.

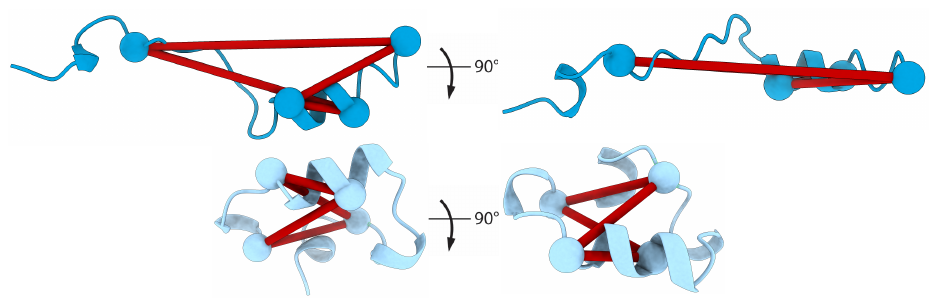

*Crosslinked residues:*

| **Residue A (within domain)** | **Residue B (within domain)** |
| --- | --- |
| 229 | 227 |
| 227 | 243 |
| 238 | 243 |
| 238 | 249 |
| **Residue A (on domain)** | **Residue B (on scaffold)** |
| 227 | 376 |
| 243 | 89 |
| 243 | 872 |
| 238 | 100 |
| 238 | 376 |
| 243 | 97 |
| 243 | 100 |
| 227 | 100 |
| 249 | 376 |
| 249 | 100 |
| 243 | 71 |
| 243 | 567 |
| 227 | 859 |
| 243 | 514 |
| 249 | 593 |
| 238 | 514 |
| 243 | 593 |
| 249 | 514 |
| 243 | 852 |
| 243 | 157 |
| 249 | 157 |
| 249 | 210 |
| 249 | 820 |

*Generated models:*

| **Model number** | **XLs /4** | **XL /23** | **QMEAN** | **z-Dope Score** | **ProCheck** | **Errat**  **Overall Quality Factor** | **ProQ2** |
| --- | --- | --- | --- | --- | --- | --- | --- |
| **ITASSER** |  |  |  |  |  |  |  |
| 1 | 4 | n/a | -2.07 | 0.44 | 3/3/2 | 58 | 15.58 |
| 2 | 4 | n/a | 0.59 | 0.49 | 2/2/4 | 50 | 9.57 |
| 3 | 4 | n/a | -1.58 | 1.13 | 4/2/3 | 54 | 13.41 |
| 4 | 4 | n/a | -3.45 | -0.19 | 4/1/3 | 58 | 9.65 |
| 5 | 4 | 19 | -2.04 | 1.23 | 3/2/3 | 58 | 15.57 |
| **Robetta msa** |  |  |  |  |  |  |  |
| 1 | 4 | n/a | 0.95 | -1.47 | 1/2/5 | 86 | 17.84 |
| 2 | 4 | n/a | 0.28 | -1.08 | 0/3/5 | 100 | 13.39 |
| 3 | 4 | n/a | 0.22 | -1.08 | 0/3/6 | 91 | 12.75 |
| 4 | 4 | n/a | 0.64 | -1.41 | 1/1/6 | 100 | 8.81 |
| **5** | **4** | n/a | **0.19** | **-1.06** | **0/0/8** | **100** | **11.73** |
| **Raptor X** |  |  |  |  |  |  |  |
| 1 | 4 | 19 | -0.61 | 1.50 | 2/2/4 | n/a | 11.93 |
| **Template-based ITASSER** |  |  |  |  |  |  |  |
| ITASSER_Model 5 as template, output 1 | 4 | 18 | -1.90 | 1.32 | 1/3/4 | 47 | 9.90 |
| ITASSER_Model 5 as template, output 2 | **4** | **20** | **-2.10** | **0.84** | **1/2/5** | **60** | **12.91** |
| Raptor X as template, output 1 | 4 | 19 | -4.29 | 1.30 | 1/4/3 | 47 | 20.53 |
| Raptor X as template, output 2 | 4 | 17 | -1.05 | 1.33 | 4/1/3 | 63 | 22.13 |

**α-Interactive Domain (α-fibrinogen’431-491)**

*Submitted Amino Acid Sequence:*

EKLVTSKGDKELRTGKEKVTSGSTTTTRRSCSKTVTKTVIGPDGHKEVTKEVVTSEDGSDC

This domain is predicted to interact to a domain from another fibrinogen chain with identical sequence. **Crosslinks can therefore not be used** for evaluation and are used only for validation after docking the domain to the scaffold. As restraint, we used preservation of an S-S bond between two highly conserved Cysteine residues. The structure of this domain was previously predicted without experimental data(2). Added to this, several in-solution NMR structures are available from previous studies of *bovine* fibrinogen and we took the most recent as template (PDB: 2jor)(3). Therefore, only ITASSER and Robetta were run in comparative mode with additional refinement for the best ITASSER model (**ITASSER, Robetta Comparative and Template-based ITASSER**). There were three close structures from Robetta and ITASSER outputs, indicating the high quality of the generated models. However, model 2 from **Robetta Comparative** was selected due to better overall scores.

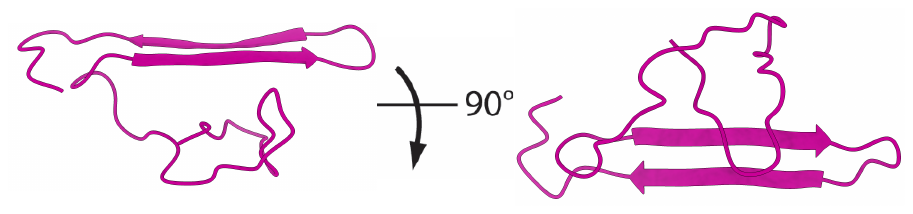

*Crosslinked residues:*

| **Residue A** | **Residue B** |
| --- | --- |
| 448 | 432 |
| 476 | 467 |
| 432 | 446 |
| 463 | 432 |
| 467 | 432 |
| 463 | 467 |
| 448 | 440 |
| 467 | 448 |
| 467 | 440 |
| 432 | 440 |
| 476 | 437 |
| 448 | 437 |
| 463 | 437 |
| 476 | 440 |
| 437 | 446 |
| 476 | 448 |
| 463 | 446 |
| 463 | 440 |
| 476 | 432 |
| 463 | 476 |

*Generated models:*

| **Model number** | **S-S**  **bond** | **XLs /20**  **(not used for evaluation)** | **ProQ2** | **QMEAN** | **z-Dope** | **ProCheck** | **Errat Overall Quality** |
| --- | --- | --- | --- | --- | --- | --- | --- |
| **ITASSER** |  |  |  |  |  |  |  |
| 1 | - | n/a | 21.57 | -7.01 | -0.22 | 4/1/3 | 39 |
| 2 | - | n/a | 29.69 | -4.29 | -0.50 | 4/1/3 | 16 |
| 3 | - | n/a | 31.38 | -4.81 | -0.42 | 4/1/3 | 50 |
| 4 | - | n/a | 31.79 | -6.38 | -0.26 | 5/1/2 | 27 |
| 5 | + | 20 | 30.94 | -3.60 | -0.31 | 2/1/5 | 21 |
| **Robetta Comparative** |  |  |  |  |  |  |  |
| 1 | - | n/a | 34.35 | -0.58 | -1.54 | 2/0/6 | 97 |
| **2** | **+** | **20** | **33.30** | **-0.26** | **-1.12** | **2/1/5** | **86** |
| 3 | + | n/a | 35.48 | -0.23 | -1.07 | 1/3/4 | 72 |
| 4 (RMSD to 2 1.185 Å) | + | 20 | 33.22 | -0.79 | -1.12 | 2/2/4 | 83 |
| 5 | + | n/a | 27.25 | 0.13 | -0.81 | 1/3/4 | 85 |
| **Template-based ITASSER** |  |  |  |  |  |  |  |
| ITASSER_Model 5 as template | - | n/a | n/a | n/a | n/a | n/a | n/a |

**α-RGD-containing Domain (α-fibrinogen’558-620)**

*Submitted Amino Acid Sequence:*

KESSSHHPGIAEFPSRGKSSSYSKQFTSSTSYNRGDSTFESKSYKMADEAGSEADHEGTHSTK

For this domain, no high quality templates were available. Even though low-confidence comparative modelling with Robetta (**Robetta Comparative**) was attempted, the models of better quality were generated by ITASSER and *ab initio* Robetta modelling (**ITASSER and Robetta msa**). In a consequent step, the best scoring ITASSER model was used as template in a refinement run, resulting in models almost identical to the template itself (**Template-based ITASSER**). Most of the models were validated by all seven restraints detected within this domain and the final selection was based on known biochemical properties. For selection we used the following rules: (1) the RGD-domain reported to interact with integrins(4) is required to be surface-exposed, and (2) the domain was hypothesized to form beta-sheets upon FXIIIa treatment which crosslinks fibrin clots(2) for which the set of residues predicted to be involved in factor XIII crosslinks are required to be surface exposed. Ultimately, this results in models where the RGD-domain is fixed between the beta-sheet. Model 5 from Robetta *ab initio* modelling was selected as it retains the factor XIII residues as surface exposed while scoring well.

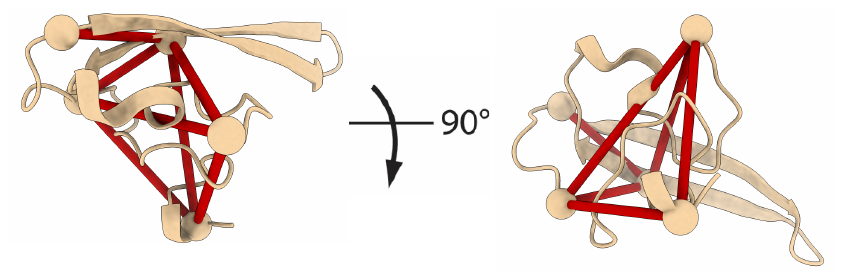

RGD domain

*Crosslinked residues:*

| **Residue A** | **Residue B** |
| --- | --- |
| 581 | 599 |
| 558 | 620 |
| 599 | 575 |
| 558 | 599 |
| 620 | 599 |
| 558 | 575 |
| 620 | 575 |

*Generated models:*

| **Model number** | **XLs/7** | **ProQ2** | **QMEAN** | **z-Dope** | **ProCheck** | **Errat Overall Quality** |
| --- | --- | --- | --- | --- | --- | --- |
| **ITASSER** |  |  |  |  |  |  |
| 1 | 7 | 31.89 | -6.89 | -0.82 | 6/3/0 | 82 |
| 2 | 7 | 31.24 | -6.34 | -0.74 | 7/1/0 | 64 |
| 3 | 7 | 29.74 | -5.45 | -1.33 | 6/3/0 | 80 |
| 4 | 7 | 28.93 | -8.97 | -1.06 | 6/3/0 | 52 |
| 5 | 7 | 35.15 | -5.89 | -1.22 | 6/2/0 | 52 |
| **Robetta msa** |  |  |  |  |  |  |
| 1 | 7 | 36.74 | 0.57 | -0.88 | 0/2/6 | 85 |
| 2 | 7 | 34.16 | 0.03 | -0.74 | 2/1/5 | 87 |
| 3 | 7 | 38.63 | 0.09 | -0.96 | 0/0/8 | 63 |
| 4 | 7 | 34.71 | 0.31 | -0.99 | 0/2/6 | 76 |
| 5 | 7 | 30.92 | 0.82 | -1.23 | 0/2/6 | 54 |
| **Robetta Comparative** |  |  |  |  |  |  |
| 1 | 7 | 34.80 | -1.03 | -0.68 | 0/2/6 | 89 |
| 2 | 7 | 40.29 | -0.99 | -0.68 | 2/1/5 | 87 |
| 3 | 5 | 24.70 | -0.52 | -0.98 | 0/1/7 | 100 |
| 4 | 7 | 29.40 | -0.66 | -1.61 | 1/1/6 | 89 |
| 5 | 7 | 26.34 | -1.42 | -0.84 | 1/2/5 | 90 |
| **Template-based ITASSER** |  |  |  |  |  |  |
| ITASSER_Model 5 as template (output is almost identical to template) | 7 | 33.62 | -7.08 | -0.71 | 5/1/2 | 2 |

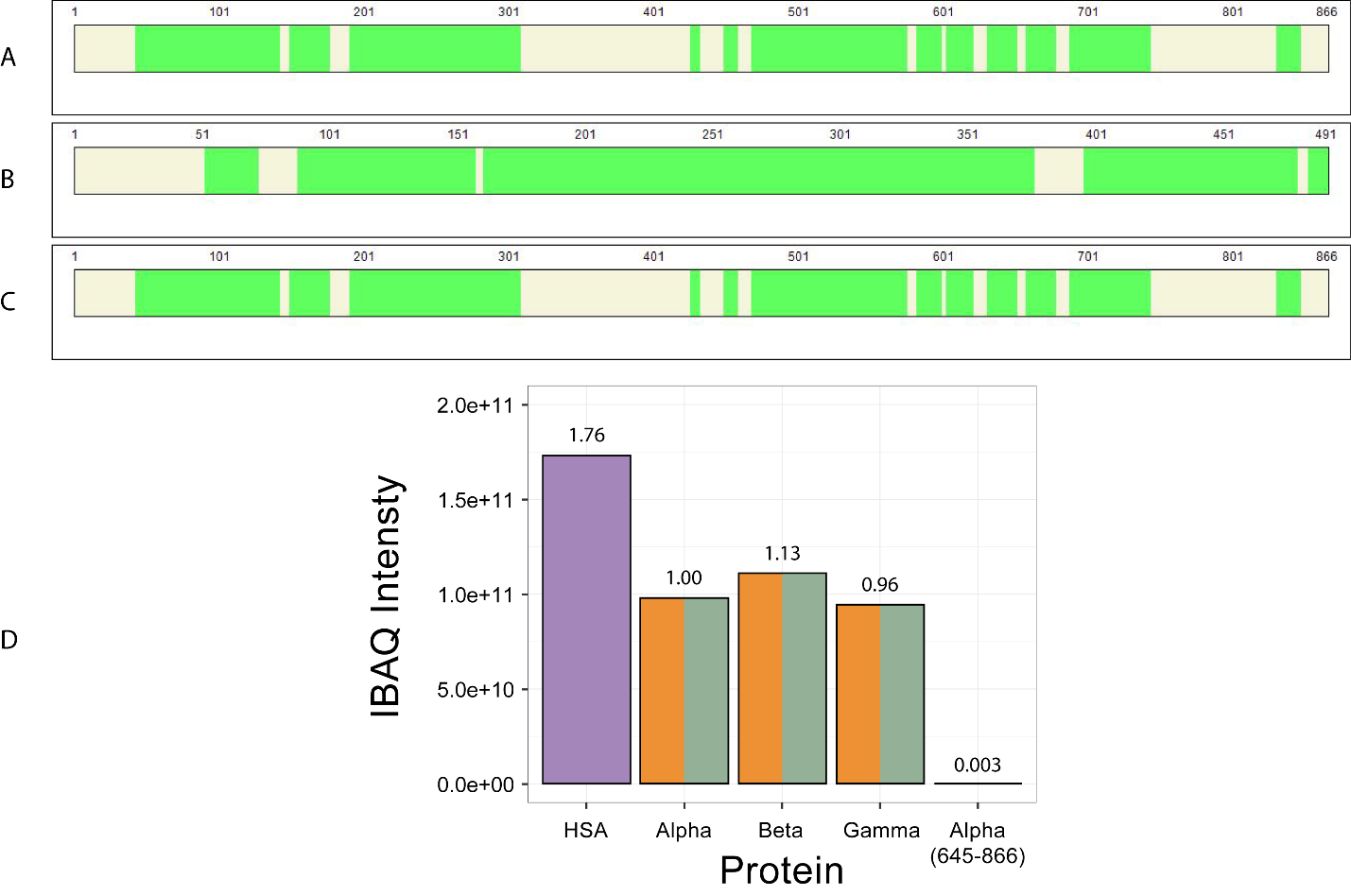

**Supplementary Figure 3:** Shotgun proteomics experiment on crosslinked fibrin clots with sequence coverage for detected peptides from shotgun mass spectrometry for **(A)** α-fibrinogen, **(B)** β-fibrinogen and **(C)** γ-fibrinogen. **(D)** Comparison of normalized intensities across a single fibrin clot unfractionated sample.

**Supplementary Note 2:** Assembling the fibrin clot based on detected distance restraints using DisVis and HADDOCK.

In all cases output was limited to 200 models.

**Assembly of the fibrin scaffold**

The available template (PDB: 3ghg) of the fibrinogen hexamer(5) was split in two trimers and submitted to CPORT for prediction of active residues (Supplementary Figure 1). Restraints identified as outlier after mapping on the template, were clustered with DisVis (Supplementary Figure 2). As we analyzed two symmetrical molecules, positions of outliers from Cluster I were swapped and analyzed again, resulting in one extra restraint for this cluster:

*Crosslinked residues:*

| **Chain** | **Residue A** | **Chain** | **Residue B** | **Cluster** |
| --- | --- | --- | --- | --- |
| Alpha | 89 | Beta | 374 | I |
| Alpha | 89 | Gamma | 406 | I |
| Alpha | 97 | Beta | 374 | I |
| Alpha | 97 | Gamma | 406 | I |
| Alpha | 100 | Beta | 374 | I |
| Alpha | 100 | Gamma | 153 | I |
| Alpha | 100 | Gamma | 406 | I |
| Alpha | 157 | Beta | 157 | I |
| Alpha | 195 | Gamma | 114 | I |
| Beta | 211 | Gamma | 114 | I |
| Beta | 295 | Alpha | 100 | I |
| Beta | 295 | Alpha | 148 | II |
| Beta | 295 | Alpha | 89 | I |
| Beta | 295 | Gamma | 114 | II |
| Beta | 313 | Alpha | 97 | I |
| Beta | 348 | Alpha | 97 | I |
| Beta | 348 | Beta | 160 | I |
| Beta | 374 | Beta | 157 | II |
| Beta | 374 | Gamma | 84 | I |
| Gamma | 61 | Beta | 374 | Outlier |
| Gamma | 114 | Gamma | 151 | I |
| Gamma | 146 | Alpha | 100 | I |
| Gamma | 146 | Gamma | 114 | I |
| Gamma | 146 | Gamma | 84 | I |
| Gamma | 153 | Alpha | 97 | I |
| Gamma | 166 | Alpha | 100 | I |
| Gamma | 399 | Gamma | 71 | I |
| Gamma | 61 | Gamma | 196 | I |
| Beta | 295 | Gamma | 121 | II |
| Beta | 295 | Gamma | 61 | Outlier |
| Beta | 374 | Gamma | 151 | Outlier |
| Beta | 374 | Alpha | 157 | II |
| Beta | 374 | Gamma | 146 | II |
| Beta | 374 | Gamma | 153 | II |
| Beta | 374 | Gamma | 177 | Outlier |
| Gamma | 299 | Beta | 374 | Outlier |

The best HADDOCK model from each cluster was selected for manual inspection. From the docking run with restraints from Cluster I, the best scoring model satisfies 18 of 24 restraints and shows no interference with expected protofibril aggregation regions (*i.e.* no interference with the central domain which is expected to be blocked by the parallel molecule). Interestingly, 5 out of 6 overlength restraints are to 2 residues on the Beta globular domain, specifically β’295 and β’374, as well as the residues involved in Cluster II. Unfortunately, docking based on these restraints did not lead to a satisfactory solution. This is likely due to the limited amount of information that can be extracted from restraints to only two positions on the large molecules involved. However, we hypothesize that these crosslinks arise from the parallel protofibril, as it is expected to be linked to exactly this region or from antiparallel alignment of β-nodules during alternative lateral aggregation of protofibrils.

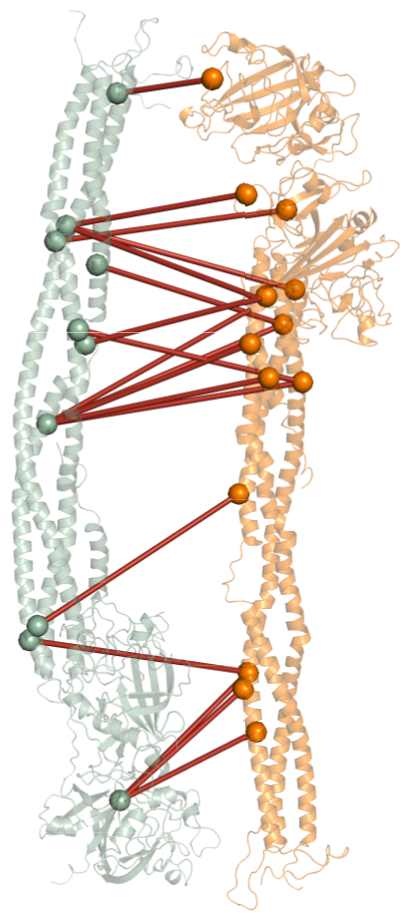

**βN-termini to Dimer**

16 out of 21 detected crosslinks between the Beta N-term and the fibrin scaffold were placed by DisVis into Cluster I (Supplementary Figure 3). However, the DisVis output indicated that all restraints are possible to map even though five outliers do not fit with most of the potential models. One of them is γ’61 and it was added to Cluster I after changing the chain to the 2^nd^ copy of Gamma. As there were potentially models satisfying all restraints, the complete set of crosslinks was submitted for docking:

*Crosslinked residues:*

| **Chain** | **Residue A** | **Chain** | **Residue B** | **Cluster** |
| --- | --- | --- | --- | --- |
| Beta | 157 | Beta | 52 | Outlier |
| Beta | 221 | Beta | 51 | Outlier |
| Beta | 295 | Beta | 51 | I |
| Beta | 295 | Beta | 52 | I |
| Beta | 348 | Beta | 52 | I |
| Beta | 353 | Beta | 52 | I |
| Beta | 367 | Beta | 51 | I |
| Beta | 374 | Beta | 51 | I |
| Beta | 374 | Beta | 77 | I |
| Beta | 422 | Beta | 51 | Outlier |
| Gamma | 61 | Beta | 51 | I |
| Gamma | 114 | Beta | 52 | Outlier |
| Gamma | 196 | Beta | 51 | I |
| Gamma | 231 | Beta | 51 | I |
| Gamma | 231 | Beta | 77 | I |
| Gamma | 231 | Beta | 83 | I |
| Gamma | 231 | Beta | 84 | I |
| Gamma | 238 | Beta | 77 | I |
| Gamma | 299 | Beta | 77 | I |
| Gamma | 299 | Beta | 83 | I |
| Gamma | 299 | Beta | 51 | I |

As the βN-terminus is known to be flexible, residues β’51-84 were defined as flexible and the best model from each output cluster was selected for manual inspection. Preservation of crosslinking restraints within the β’N-termini was required (Supplementary Note 1) and only models satisfying this criterion was selected for further analysis. From all available models, the best satisfies a total of 16 restraints. As the remaining crosslinks involve residues β’51 and β’52 located on the β’N-termini, these were excluded as placement of this flexible domain cannot effectively be verified with crosslinks.

**α-Elongated and α-Folded Domains to Dimer**

As these domains were modelled as an extension of the existing template, we fused the predicted models to the available structure. Placement was done according to the detected crosslinks (Supplementary Note 1) to the assembled scaffold. To correct for inaccuracies introduced by manual placement, HADDOCK in Refinement mode was used. The top scoring models from the refinement output were selected for further assembly steps. Further docking steps were performed mostly on the region of the molecule containing the α-Folded conformation. The resulting models from each of the following steps were confirmed not to interfere with newly docked domains. Eventually, all the domains described below are equally fitting to the α-Elongated and α-Folded conformations. On the final fibrin clot model, mirrored versions of both domains were examined to obtain a full picture of potentially important mutation sites.

**α-Interactive Domains to Dimer**

This domain has the largest number of detected inter-molecular crosslinks. A total of 69 restraints were clustered with DisVis (Supplementary Figure 4):

*Crosslinked residues:*

| **Chain** | **Residue A** | **Chain** | **Residue B** | **Cluster** |
| --- | --- | --- | --- | --- |
| Beta | 52 | Alpha | 440 | I |
| Beta | 77 | Alpha | 437 | I |
| Beta | 83 | Alpha | 437 | I |
| Alpha | 157 | Alpha | 467 | I |
| Alpha | 157 | Alpha | 476 | I |
| Alpha | 157 | Alpha | 432 | I |
| Alpha | 157 | Alpha | 437 | I |
| Alpha | 157 | Alpha | 440 | I |
| Alpha | 157 | Alpha | 446 | I |
| Alpha | 157 | Alpha | 448 | I |
| Alpha | 202 | Alpha | 463 | I |
| Alpha | 202 | Alpha | 432 | I |
| Alpha | 202 | Alpha | 437 | I |
| Alpha | 202 | Alpha | 440 | I |
| Alpha | 202 | Alpha | 476 | I |
| Alpha | 210 | Alpha | 440 | I |
| Alpha | 210 | Alpha | 448 | I |
| Beta | 295 | Alpha | 437 | I |
| Beta | 295 | Alpha | 440 | I |
| Beta | 295 | Alpha | 432 | I |
| Beta | 295 | Alpha | 448 | I |
| Beta | 295 | Alpha | 446 | I |
| Beta | 313 | Alpha | 440 | I |
| Beta | 348 | Alpha | 437 | I |
| Beta | 348 | Alpha | 440 | I |
| Beta | 348 | Alpha | 467 | I |
| Beta | 348 | Alpha | 476 | I |
| Beta | 348 | Alpha | 432 | I |
| Beta | 348 | Alpha | 463 | I |
| Beta | 353 | Alpha | 440 | I |
| Beta | 367 | Alpha | 440 | I |
| Beta | 374 | Alpha | 437 | I |
| Beta | 374 | Alpha | 440 | I |
| Beta | 374 | Alpha | 467 | I |
| Beta | 374 | Alpha | 448 | I |
| Beta | 374 | Alpha | 446 | I |
| Beta | 374 | Alpha | 476 | I |
| Beta | 422 | Alpha | 467 | I |
| Beta | 422 | Alpha | 432 | I |
| Beta | 426 | Alpha | 440 | I |
| Beta | 426 | Alpha | 432 | I |
| Gamma | 146 | Alpha | 432 | I |
| Gamma | 151 | Alpha | 437 | I |
| Gamma | 151 | Alpha | 440 | I |
| Gamma | 151 | Alpha | 463 | I |
| Gamma | 166 | Alpha | 440 | I |
| Beta | 51 | Alpha | 432 | II |
| Beta | 51 | Alpha | 437 | II |
| Beta | 51 | Alpha | 440 | II |
| Beta | 71 | Alpha | 448 | II |
| Alpha | 89 | Alpha | 432 | II |
| Alpha | 89 | Alpha | 476 | II |
| Alpha | 89 | Alpha | 440 | II |
| Alpha | 97 | Alpha | 448 | II |
| Alpha | 100 | Alpha | 440 | II |
| Alpha | 100 | Alpha | 448 | II |
| Beta | 295 | Alpha | 467 | II |
| Beta | 295 | Alpha | 463 | II |
| Beta | 367 | Alpha | 463 | II |
| Beta | 367 | Alpha | 476 | II |
| Gamma | 61 | Alpha | 432 | II |
| Gamma | 61 | Alpha | 440 | II |
| Gamma | 146 | Alpha | 448 | II |
| Gamma | 166 | Alpha | 448 | II |
| Gamma | 177 | Alpha | 440 | II |
| Gamma | 114 | Alpha | 476 | III |
| Gamma | 114 | Alpha | 437 | III |
| Gamma | 114 | Alpha | 440 | III |
| Gamma | 114 | Alpha | 432 | III |

In all docking runs, parts of the interactive alpha domain outside the beta-sheet are defined as flexible (residues α432-460).

**Cluster I** HADDOCK docking resulted in only 46 possible structures. From the top models of each cluster, we performed manual validation and selected the model satisfying the largest number of restraints.

**Cluster II** was HADDOCK docked to the model with Cluster I in place and resulted in only 59 possible structures. From the top models of each cluster, we performed manual validation and selected the model satisfying the largest number of restraints. In this case, all 19 crosslinks used for docking were within the set distance constraint. Two outliers from the Cluster I docking run were mapped on the structure obtained from docking with Cluster II and remain outside the maximum allowed distance constraint.

**Cluster III** was HADDOCK docked to the model with Cluster I and Cluster II in place. From the top model of each cluster, we performed manual validation and selected the model which satisfies the largest number of restraints. The α-domain was placed in the middle of the coiled-coil region. This may be explained either by a transitional state towards the active form. However, due to the limited amount of crosslinks all focused on a single residue this model was excluded.

**α-RGD-containing Domains to Dimer**

21 out of 33 detected crosslinks between the α-RGD-containing Domain and the Fibrin Scaffold were placed by DisVis in Cluster I (Supplementary Figure 5):

*Crosslinked residues:*

| **Chain** | **Residue A** | **Chain** | **Residue B** | **Cluster** |
| --- | --- | --- | --- | --- |
| Alpha | 157 | Alpha | 558 | I |
| Alpha | 157 | Alpha | 575 | I |
| Alpha | 157 | Alpha | 599 | I |
| Alpha | 157 | Alpha | 581 | I |
| Alpha | 157 | Alpha | 620 | I |
| Alpha | 195 | Alpha | 558 | I |
| Alpha | 210 | Alpha | 599 | I |
| Alpha | 210 | Alpha | 558 | I |
| Alpha | 210 | Alpha | 575 | I |
| Beta | 52 | Alpha | 575 | I |
| Beta | 295 | Alpha | 620 | I |
| Beta | 295 | Alpha | 558 | I |
| Beta | 295 | Alpha | 575 | I |
| Beta | 295 | Alpha | 581 | I |
| Beta | 295 | Alpha | 599 | I |
| Beta | 367 | Alpha | 575 | I |
| Beta | 367 | Alpha | 599 | I |
| Gamma | 146 | Alpha | 575 | I |
| Gamma | 151 | Alpha | 575 | I |
| Gamma | 151 | Alpha | 599 | I |
| Gamma | 151 | Alpha | 620 | I |
| Alpha | 100 | Alpha | 558 | II |
| Beta | 77 | Alpha | 575 | II |
| Beta | 348 | Alpha | 558 | II |
| Beta | 348 | Alpha | 620 | II |
| Beta | 348 | Alpha | 575 | II |
| Beta | 353 | Alpha | 620 | II |
| Beta | 374 | Alpha | 558 | II |
| Beta | 374 | Alpha | 599 | II |
| Gamma | 114 | Alpha | 599 | II |
| Gamma | 114 | Alpha | 575 | II |
| Gamma | 166 | Alpha | 575 | II |
| Gamma | 177 | Alpha | 620 | II |

It is expected that parts of this domain apart from the modelled beta-sheet are flexible and are defined in HADDOCK accordingly (α558-580, α604-620).

**Cluster I** HADDOCK docking resulted in only 28 possible structures. The best models from each generated cluster were inspected manually. On top of the number of validated restraints, it was enforced that integrin-binding RGD-domain is exposed. The selected model satisfies these criteria and 18 restraints.

**Cluster II** was HADDOCK docked to a model with Cluster I in place and resulted in only 14 possible structures. We examined the best models manually. On top of the number of validated restraints, also here it was enforced that the integrin-binding RGD-domain is exposed. The selected model satisfies these criteria and seven restraints.

Outliers from both clusters were mapped on an alternative placement of the domain. This lead to an extra two validated restraints with six outliers left in total.

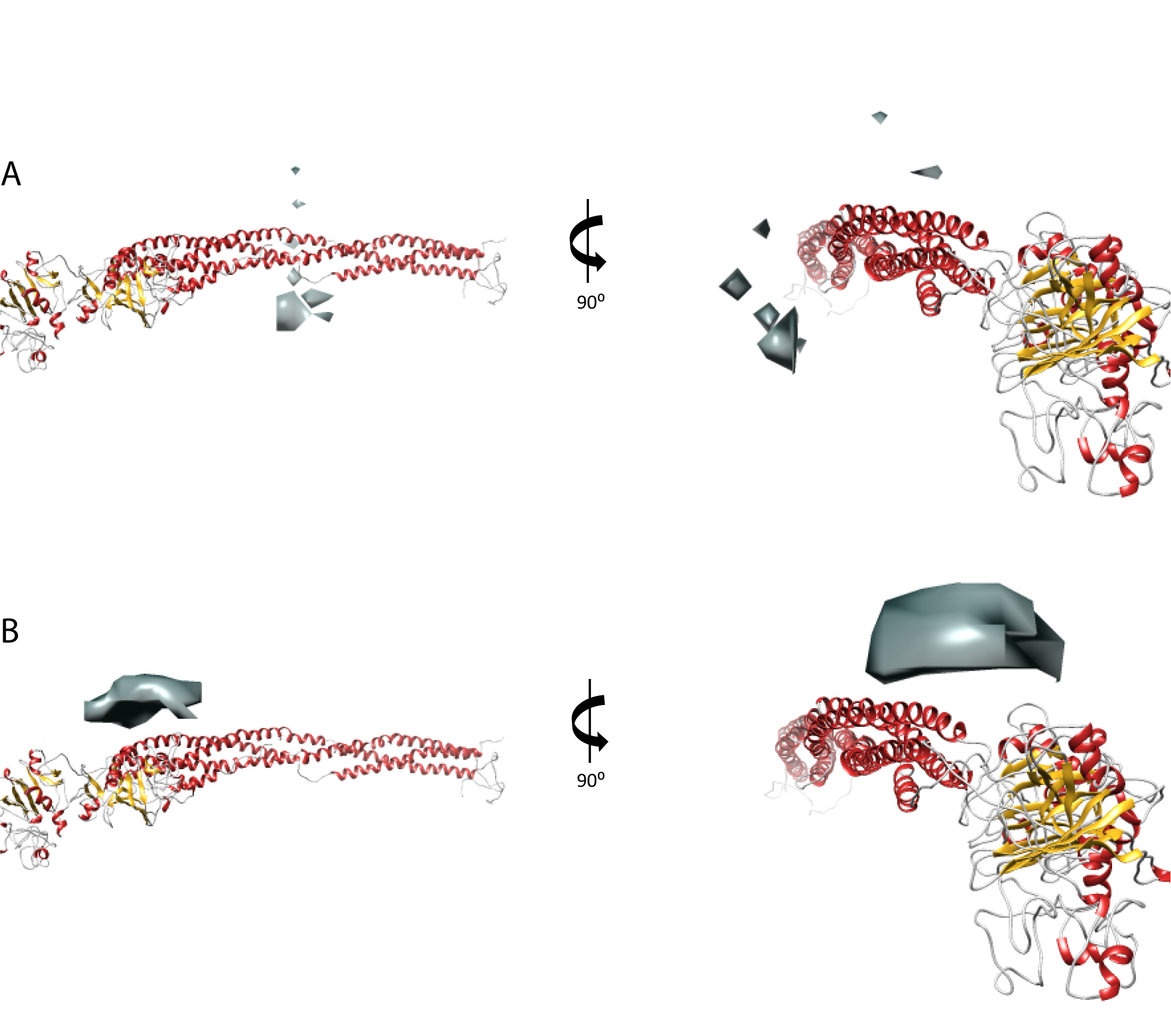

**Supplementary Figure 4:** DisVis analysis of the interaction interface between the fibrinogen chains based on the overlength restraints after mapping onto the known template (PDB: 3ghg). **(A)** Cluster I with the predicted interface (grey) for 16 restraints and **(B)** Cluster II with the predicted interface (grey) for six restraints. The number of restraints is adjusted to make the interaction interface visible.

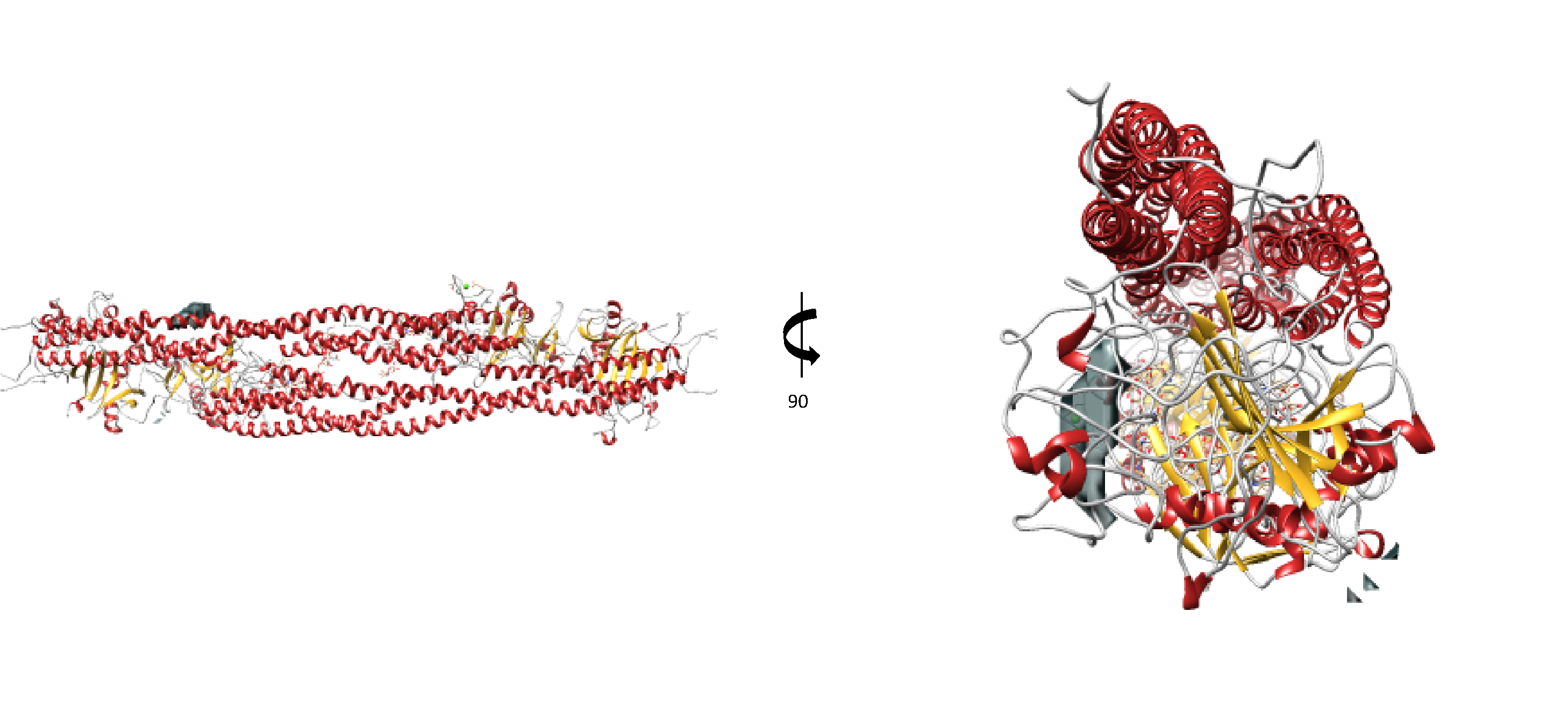

**Supplementary Figure 5:** DisVis analysis of the interaction interface βN-terminal domain and fibrin based on the detected restraints. All 21 submitted crosslinks are found to be ‘True Positives’ and grouped into a single cluster.

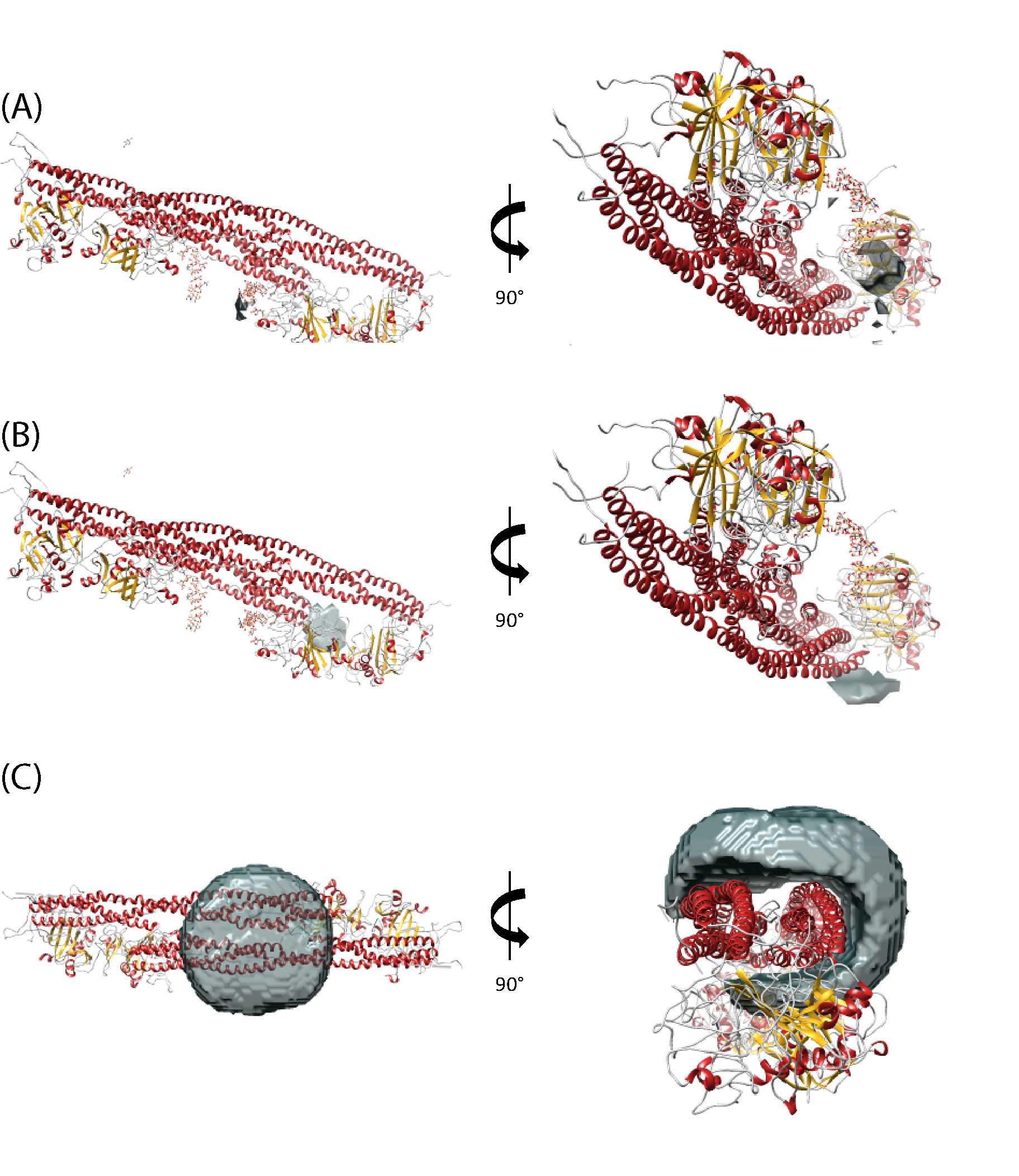

**Supplementary Figure 6:** DisVis analysis of the interaction interface between αN-terminal sub-domain and fibrin based on detected restraints. **(A)** Cluster I with the predicted interface (grey) for 42 restraints. **(B)** Cluster II with the predicted interface (grey) for 16 restraints. The number of restraints is adjusted to make the interaction interface visible. **(C)** Cluster III with the predicted interface (grey) for four restraints. The number of restraints is adjusted to make the interaction interface visible.

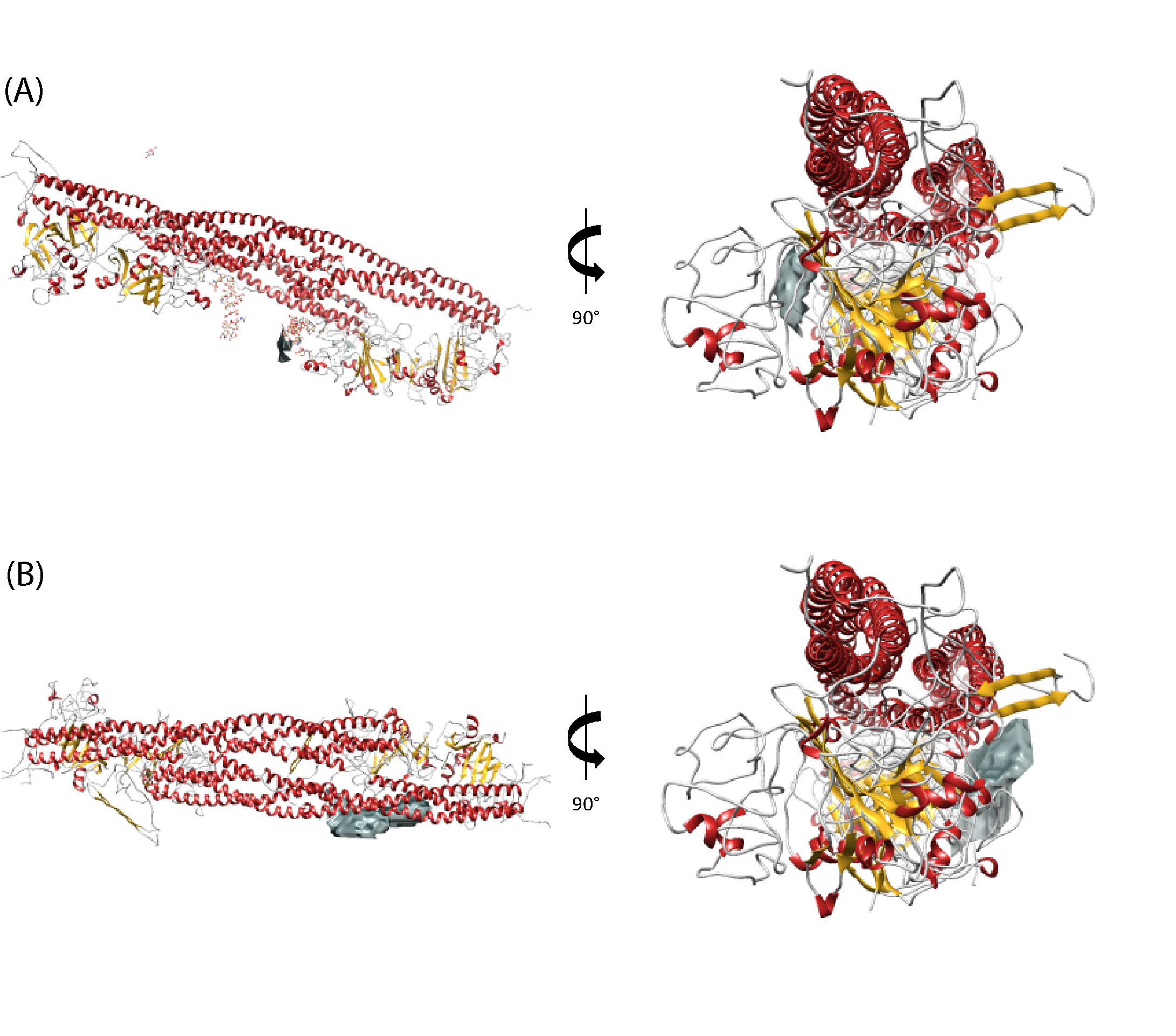

**Supplementary Figure 7:** DisVis analysis of the interaction interface between αC-terminal subdomain and fibrin based on the detected restraints. **(A)** Cluster I with the predicted interface (grey) is relatively large and is not strictly defined even for 40 restraints. Nevertheless, only 21 restraint have been grouped into a cluster. **(B)** Cluster II with the predicted interface (grey) for nine restraints. The number of restraints is adjusted to make the interaction interface visible.

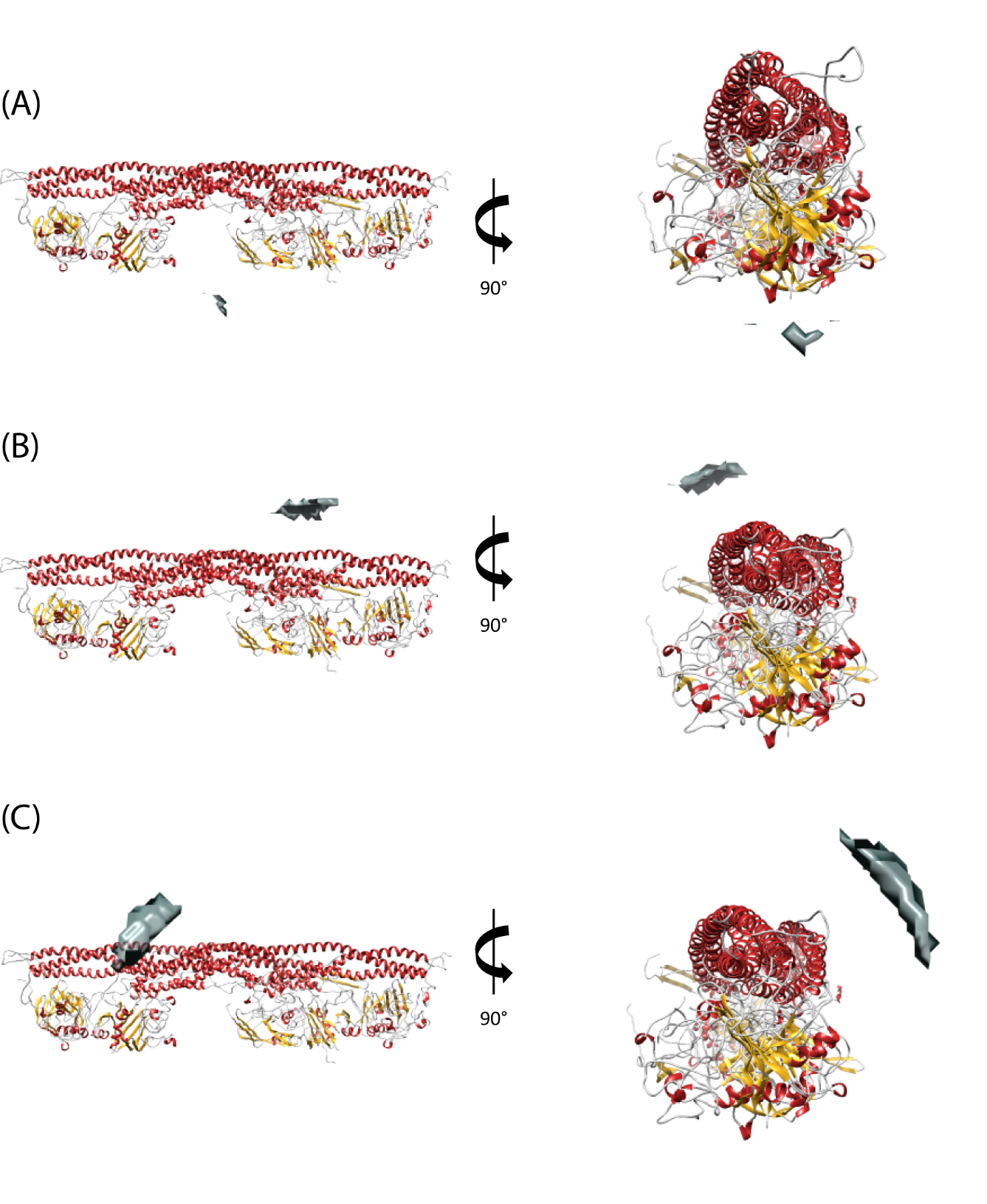

**Supplementary Figure 8:** DisVis analysis of HSA to fibrin. The number of restraints is adjusted to make the interaction interface visible.

**Supplementary Table 3:** Mapping of the detected intermolecular restraints between HSA (Uniprot accession code P02678) and fibrin clot (Uniprot accession codes P02671 for α-Fibrinogen and P02675 for β-Fibrinogen) on the generated models.

| **Sequence A** | **Accession A** | **Position A** | **Sequence B** | **Accession B** | **Position B** | **Distance, Å** | **Cluster I** | **Cluster II** |
| --- | --- | --- | --- | --- | --- | --- | --- | --- |
| NN[K]DSHSLTTNIMEILR | P02671 | 100 | [K]YLYEIAR | P02768 | 161 |  | - | 23.48 |
| VQHIQLLQ[K]NVR | P02671 | 157 | F[K]DLGEENFK | P02768 | 36 |  | 39.94 | 34.49 |
| VQHIQLLQ[K]NVR | P02671 | 157 | [K]YLYEIAR | P02768 | 161 |  | - | 41.12 |
| VQHIQLLQ[K]NVR | P02671 | 157 | ETCFAEEG[K]K | P02768 | 597 |  | - | - |
| SQLQ[K]VPPEWK | P02671 | 243 | [K]YLYEIAR | P02768 | 161 |  | 38.10 | - |
| EYHTE[K]LVTSK | P02671 | 432 | [K]YLYEIAR | P02768 | 161 |  | 31.64 | 26.57 |
| TVT[K]TVIGPDGHK | P02671 | 467 | [K]YLYEIAR | P02768 | 161 |  | 29.70 | 41.53 |
| TVIGPDGH[K]EVTK | P02671 | 476 | [K]YLYEIAR | P02768 | 161 |  | 35.34 | - |
| GDSTFES[K]SYK | P02671 | 599 | [K]YLYEIAR | P02768 | 161 |  | 43.31 | - |
| YQISVN[K]YR | P02675 | 374 | F[K]DLGEENFK | P02768 | 36 |  | - | - |
| YQISVN[K]YR | P02675 | 374 | SLHTLFGD[K]LCTVATLR | P02768 | 97 |  | - | - |
| [K]WDPYK | P02675 | 295 | SLHTLFGD[K]LCTVATLR | P02768 | 97 |  | 34.91 | - |
| [K]REEAPSLRPAPPPISGGGYR | P02675 | 52 | [K]YLYEIAR | P02768 | 161 |  | - | 31.32 |
| YQISVN[K]YR | P02675 | 374 | [K]YLYEIAR | P02768 | 161 |  | 39.44 | - |
| YQISVN[K]YR | P02675 | 374 | AAFTECCQAAD[K]AACLLPK | P02768 | 198 |  | 36.39 | - |
| YQISVN[K]YR | P02675 | 374 | AF[K]AWAVAR | P02768 | 236 |  | 42.91 | - |

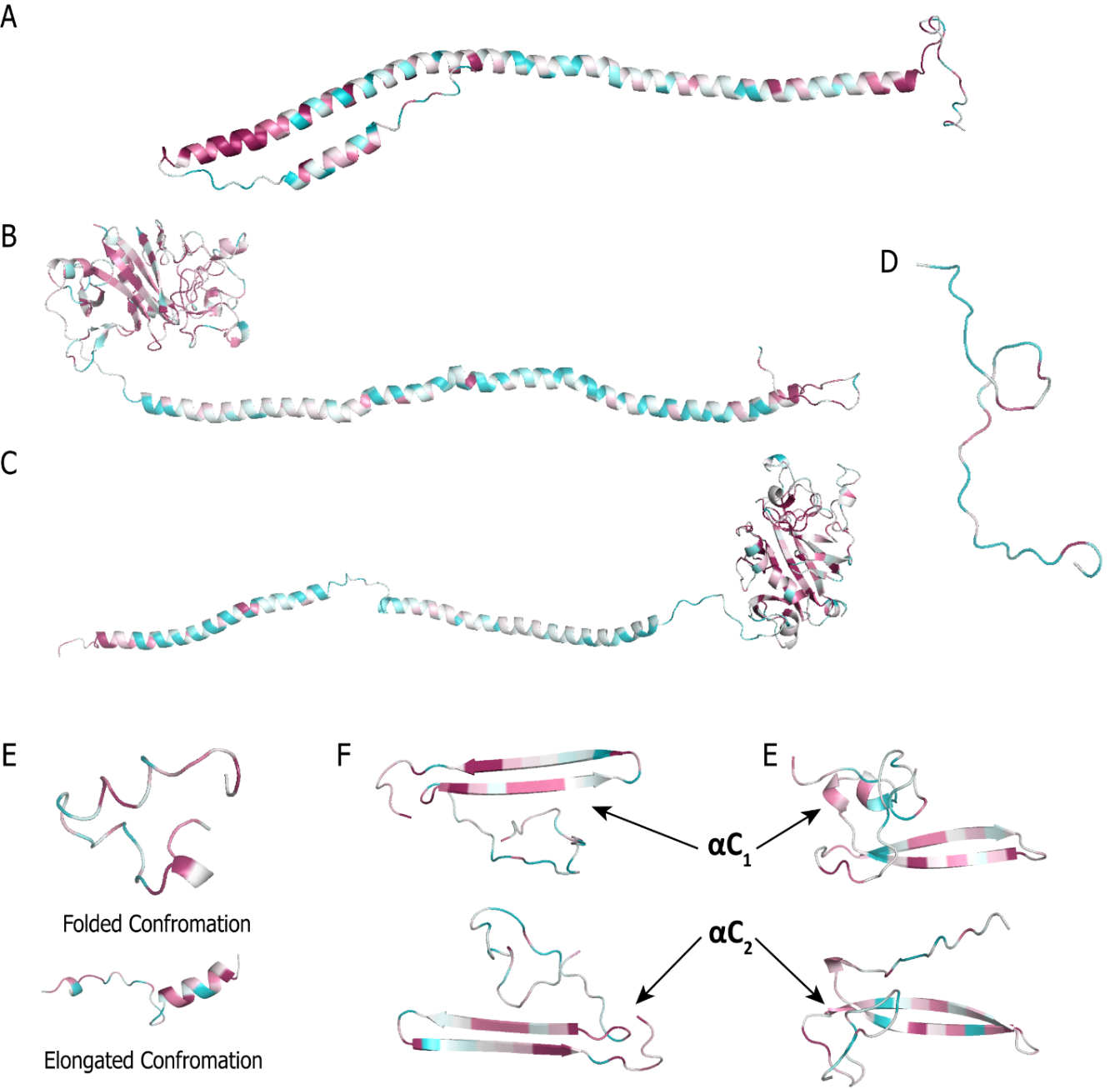

**Supplementary Data 1.** Conservation Analysis of sequences from fibrin. Conservation of residues varies from green (low scores) through white (moderate scores) to dark red (high scores). **(A)** The α-chain from template PDB: 3ghg (α-fibrinogen’46-219). **(B)** The β-chain from template PDB: 3ghg (Fibrinogen β’88-458). **(C)** The γ-chain form template PDB: 3ghg (Fibrinogen γ’40-420). **(D)** The βN-term (Fibrinogen β’51-84). **(E)** Two conformations of the modelled α-chain domain (α-fibrinogen’220-249). **(F)** Both α-Interactive Domains (α-fibrinogen’431-491) after docking onto the final fibrin clot structural model. **(G)** Both α-RGD-containing domains (α-fibrinogen’558-620) from the final fibrin clot model.

**Supplementary Data 2-8.** Consurf color-coded Multiple Sequence Alignments for each residue of fibrin clots including relevant sequences.

See PDF file ‘Supplementary Data 2-8.pdf’:

**Supplementary Data 2:** α-chain from template 3ghg (α-fibrinogen’46-219) **p.1**

**Supplementary Data 3:** 1st α-chain Domain (α-fibrinogen’220-249) **p.14**

**Supplementary Data 4:** α-Interactive Domains (α-fibrinogen’431-491) **p.16**

**Supplementary Data 5**: α-RGD-containing Domains (α-fibrinogen’558-620) **p.20**

**Supplementary Data 6:** βN-term (Fibrinogen β’51-84) **p.23**

**Supplementary Data 7:** β-chain from template 3ghg (Fibrinogen β’88-458) **p.25**

**Supplementary Data 8:** γ-chain from template 3ghg (Fibrinogen γ’40-420) **p.102**

**Supplementary Movie:** Movie explaining the advances made with this study

**Supplementary Table 4:** Full list of mapped relevant mutations.

| **Gene** | **Amino acid change** | **Associated to / source** | **CADD score** | **Conservation score** | **Potential effect** | **Conservation scores of affected residues** |
| --- | --- | --- | --- | --- | --- | --- |
| **FGA** | S453N | Maekawa et. al.(6)  The model provides an additional function | 21.6 | 1 | Extra glycosylation might interfere with the correct folding and placement of the α432-491 domain. | - |
|  | C491S | Ikeda et. al.(7) | 27.6 | 9 | Lack of the interactive beta-sheet in the α432-491 domain. Leaves free Cys’461 which forms a complex with albumin. | 9 |
|  | E559V | Gillmore et. al.(8) | 13.9 | - | Change of charged Glu to non-polar Val can significantly affect the flexibility of that region. Rotamer gives a salt bridge to αLys'157, *i.e.* important in the placement of α558-620 with RGD-domain. | 2 |
|  | P571H | Amyloidosis(9) | 17.0 | - | Change of hydrophobic Pro’571 to polar His - affects flexibility through *e.g.* π-π stacking with Phe'570 or cation-π interaction with Arg'573.  The His gives rise to a H-π interaction with Ser'576 and interferes with its function implied by its high conservation score.  Alternatively, H-π with Glu’240 is also possible. Another rotamer provides the potential for π-π stacking with Tyr'601, fixing an otherwise flexible loop into a beta-sheet extension. | αPhe’570-1  αSer’576-9  αTyr’601-1 |
|  | R573C R573H, R573L | Dysfibrinogenemia(10, 11), Amyloidosis(12) | 18.1, 13.4, 13.7 | 6 | In αC_1_, αArg’573 is involved in an H-H bond with the side chain of αLys'225 in the folded conformation. In other organisms often changed to Ser or Thr, also capable of maintaining this interaction. Within αC_2_, has the potential to form a salt bridge with αGlu’447.  Change to His provides the potential for π-π stacking to the neighboring αPhe’570 and potentially a cation-π interaction to αLys'227.  Amyloidosis-causing substitution to Leu provides the potential for α580-603 beta-sheet extension. | αLys’225-9  αGlu’447-6;  αPhe’570-1;  αLys’227-1 |
|  | G574F | Rowczenio et.al.(13) | - | 6 | Change of Gly to Phe, potentially involved in a π-cation with Arg'573 or π-π stacking with Phe'570. Also, in our model disrupts π-π stacking to Tyr’601. | 1 |
|  | M603L | Schönland et. al.(14) | 7.9 | 5 | Change to Leu may force a turn of the flexible domain and cause an extension of the beta-sheets α580-603. | - |
| **FGB** | ΔD39-L102 | Liu et. al.(15) | - | - | Thrombotic tendency due to lack of the beta knob, *e.g.* no "opener" for the α148-160 plasminogen-binding site. | - |
|  | Y71N | Marchi et. al.(16) | 26.2 | 9 | Close proximity to βArg'72 and βArg'74 within the Heparin Binding Domain. Part of the loop, which can be extended and inserts knob b into hole B. | - |
|  | R74C, R74G, R74L | Koopman et. al. (17); Würtinger et. al.(18); Shlebak et.al.(19)  Extra salt bridge from our model | 34.0; 29.1; 31.0 | 9 | Change of a highly conserved Arg, which is close to FbB cleavage site, affects the flexibility of β51-83. In our model, involved in a salt bridge to βAsp'350. Explains thrombosis tendency through the disturbed “opener” of t-PA α148-160 domain. | 4 |
|  | Q118* | Casini et.al.(20) | 31.0 | 4 | Lack of a structurally functional knob B and βArg’196. | - |
|  | ΔSer141 | Okumura et. al.(21);  The potential function is mentioned, but our model provides evidence | - | 7 | Shifts of the coiled-coil residues. The residue itself does not affect lateral aggregation. | - |
|  | M148K | Predicted (CADD). Region β148-164 is mentioned in Lugovskoy et. al.(22) | 6.2 | 4 | Helix disruption, potentially has an effect on the salt bridge of βArg’196. | - |
|  | N190S | Sugo et. al.(23);  The model provides an additional function | 0.1 | 2 | Extra glycosylation is in close proximity to parallel protofibril, would interfere with lateral aggregation. Also, βN'188 forms H-H bond to βE'185. | 6 |
|  | L195P | Mitchell (2002) FGB LSDB Unpublished | 23.8 | 8 | In close proximity to βArg’196. | - |
|  | R196C | Lounes et. al.(24); Dysfibrinogenemia | 28.9 | 6 | In addition to the formation of albumin-complexes, βArg’196 is crucial for lateral aggregation through a salt bridge with βGlu’177. In some other organisms, this residue is changed to Lys. | βGlu’177-8 |
|  | R478K | Ajjan et. al.(9);  The model provides an alternative hypothesis | 0.1 | 8 | Guiding residue for Beta-knob. Potential H-H bond between Arg’478 side-chain to the backbone of βPro57, which would be not possible with Lys. | 2 |
| **FGG** | Y140H | Morris et. al.(25);  The model provides an additional function | 22.0 | 5 | Polar imidazole within the ‘helix-permissive’ hydrophobic center. In our model, this helix disruption affects the region involved in lateral aggregation. | - |

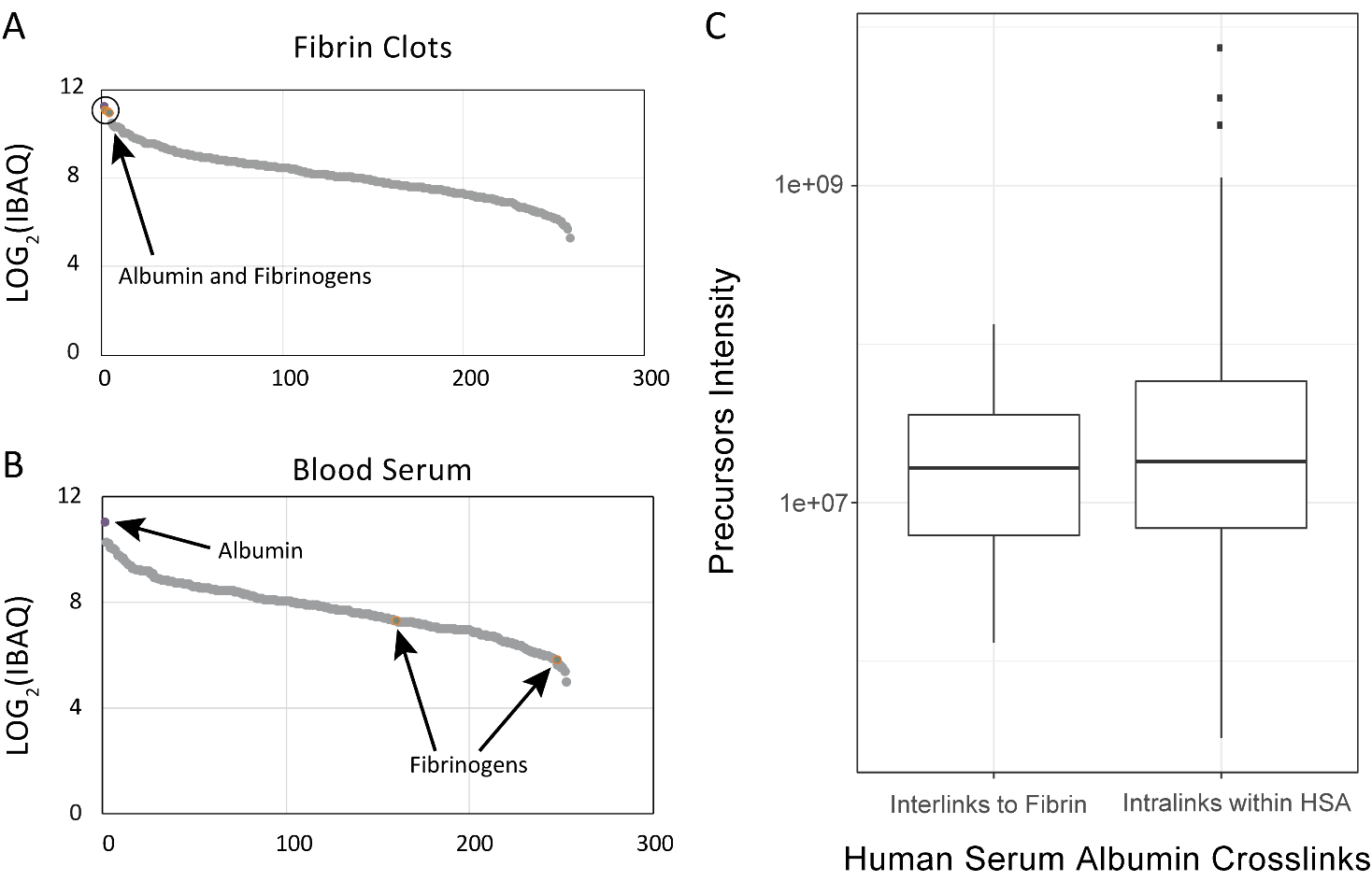
**Supplementary Figure 9.** Specificity of albumin interaction with fibrin clots. **(A, B)** S-curves of log_2_(iBAQ) values for proteins in purified clots and blood serum; obtained with MaxQuant **i**ntensity-**B**ased **A**bsolute **Q**uantification this indicates a very defined stoichiometry. **(C)** Comparison of precursor intensity of crosslinked peptides. On the left, the intensity of crosslinks between albumin and fibrin clots. On the right, intensity of crosslinks within albumin. Average intensities of precursors are almost identical in both cases, indicating a strong interaction.

**Supplementary Note 3:** Deciphering the interaction interface between fibrin and HSA based on detected distance restraints.

Docking of slbumin to fibrin clots was performed with **ClusPro** in restraints mode. In all cases, 50% of defined restraints were excluded. Models were selected on **balanced scores** and detected crosslinks.

**Clustering of restraints with DisVis**

There are 16 restraints in the final dataset between albumin and the fibrin clot (**Supplementary Table 3**). However, as two copies of each fibrinogen subunit exist in the fibrin clot structure, 15 of the retrains can belong to 2 different chains. In the case of albumin, only monomers are expected to be present.

| **Accession A**  **(Fibrinogens)** | **Chains in PDB-DEV**  **(Fibrinogens)** | **Position A**  **(Fibrinogens)** | **Accession B (Albumin)** | **Position B** |
| --- | --- | --- | --- | --- |
| P02671 | A, D | 100 | P02768 | 161 |
| P02671 | A, D | 157 | P02768 | 36 |
| P02671 | A, D | 157 | P02768 | 161 |
| P02671 | A, D | 157 | P02768 | 597 |
| P02671 | Y, Z | 243 | P02768 | 161 |
| P02671 | G, H | 432 | P02768 | 161 |
| P02671 | G, H | 467 | P02768 | 161 |
| P02671 | G, H | 476 | P02768 | 161 |
| P02671 | I, J | 599 | P02768 | 161 |
| P02675 | B, E | 374 | P02768 | 36 |
| P02675 | B, E | 374 | P02768 | 97 |
| P02675 | B, E | 295 | P02768 | 97 |
| P02675 | K | 52 | P02768 | 161 |
| P02675 | B, E | 374 | P02768 | 161 |
| P02675 | B, E | 374 | P02768 | 198 |
| P02675 | B, E | 374 | P02768 | 236 |

Eventually, 16 crosslinks result in 31 possible restraints between each copy of fibrinogen and albumin. All possible restraints were clustered with DisVis onto the structure of the fibrin clot using the available albumin structure (PDB: 1uor):

| **Chain in PDB-DEV**  **(Fibrinogens)** | **Residue A (Fibrinogens)** | **Residue B**  **(Albumin)** | **Cluster** |
| --- | --- | --- | --- |
| A | 100 | 161 | III  (Copied in II) |
| A | 157 | 36 | II |
| A | 157 | 161 | II |
| A | 157 | 597 | II |
| Y | 243 | 161 | I |
| G | 432 | 161 | I |
| G | 467 | 161 | I |
| G | 476 | 161 | I |
| I | 599 | 161 | I |
| B | 374 | 36 | I |
| B | 374 | 97 | I |
| B | 295 | 97 | II |
| K | 52 | 161 | II |
| B | 374 | 161 | I |
| B | 374 | 198 | I |
| B | 374 | 236 | Outlier  (copied in I) |
| D | 100 | 161 | II |
| D | 157 | 36 | I |
| D | 157 | 161 | III  (Copied in II) |
| D | 157 | 597 | III  (Copied in II) |
| D | 243 | 161 | I |
| H | 432 | 161 | II |
| H | 467 | 161 | II |
| H | 476 | 161 | II |
| J | 599 | 161 | I |
| E | 374 | 36 | I |
| E | 374 | 97 | I |
| E | 295 | 97 | III  (Copied in II) |
| E | 374 | 161 | I |
| E | 374 | 198 | I |
| E | 374 | 236 | I |

**Cluster I:** Docking with defined restraints resulted in 764 models. Outputs were clustered on Balanced score:

| **Cluster** | **Members** | **Representative** | **Weighted Score** |
| --- | --- | --- | --- |
| **0** | 63 | Center | -711.69 |
|  |  | Lowest Energy | -965.00 |
| **1** | 54 | Center | -698.24 |
|  |  | Lowest Energy | -999.46 |
| **2** | 41 | Center | -827.60 |
|  |  | Lowest Energy | -1002.39 |
| **3** | 41 | Center | -743.37 |
|  |  | Lowest Energy | -956.89 |
| **4** | 37 | Center | -704.58 |
|  |  | Lowest Energy | -916.76 |
| **5** | 36 | Center | -655.37 |
|  |  | Lowest Energy | -745.36 |
| **6** | 31 | Center | -760.39 |
|  |  | Lowest Energy | -833.00 |
| **7** | 30 | Center | -729.14 |
|  |  | Lowest Energy | -845.22 |
| **8** | 29 | Center | -720.57 |
|  |  | Lowest Energy | -954.66 |
| **9** | 29 | Center | -725.32 |
|  |  | Lowest Energy | -891.02 |
| **10** | 28 | Center | -694.48 |
|  |  | Lowest Energy | -779.58 |
| **11** | 27 | Center | -677.70 |
|  |  | Lowest Energy | -1097.14 |
| **12** | 27 | Center | -681.72 |
|  |  | Lowest Energy | -765.19 |
| **13** | 26 | Center | -750.14 |
|  |  | Lowest Energy | -978.68 |
| **14** | 24 | Center | -790.54 |
|  |  | Lowest Energy | -914.58 |
| **15** | 24 | Center | -681.42 |
|  |  | Lowest Energy | -777.19 |
| **16** | 22 | Center | -707.54 |
|  |  | Lowest Energy | -796.45 |
| **17** | 21 | Center | -730.34 |
|  |  | Lowest Energy | -1047.81 |
| **18** | 20 | Center | -664.03 |
|  |  | Lowest Energy | -977.25 |
| **19** | 16 | Center | -722.79 |
|  |  | Lowest Energy | -810.12 |
| **20** | 16 | Center | -675.80 |
|  |  | Lowest Energy | -881.46 |
| **21** | 16 | Center | -652.13 |
|  |  | Lowest Energy | -765.76 |
| **22** | 16 | Center | -695.97 |
|  |  | Lowest Energy | -726.92 |
| **23** | 15 | Center | -670.37 |
|  |  | Lowest Energy | -947.77 |
| **24** | 15 | Center | -751.09 |
|  |  | Lowest Energy | -751.09 |
| **25** | 13 | Center | -700.20 |
|  |  | Lowest Energy | -777.00 |
| **26** | 12 | Center | -714.53 |
|  |  | Lowest Energy | -860.16 |
| **27** | 12 | Center | -726.47 |
|  |  | Lowest Energy | -726.47 |
| **28** | 12 | Center | -733.98 |
|  |  | Lowest Energy | -802.01 |
| **29** | 11 | Center | -813.54 |
|  |  | Lowest Energy | -813.54 |

We evaluated five models manually, and eventually the 3^rd^-best model was selected as it satisfies the most restaints (11 restraints out of 17).

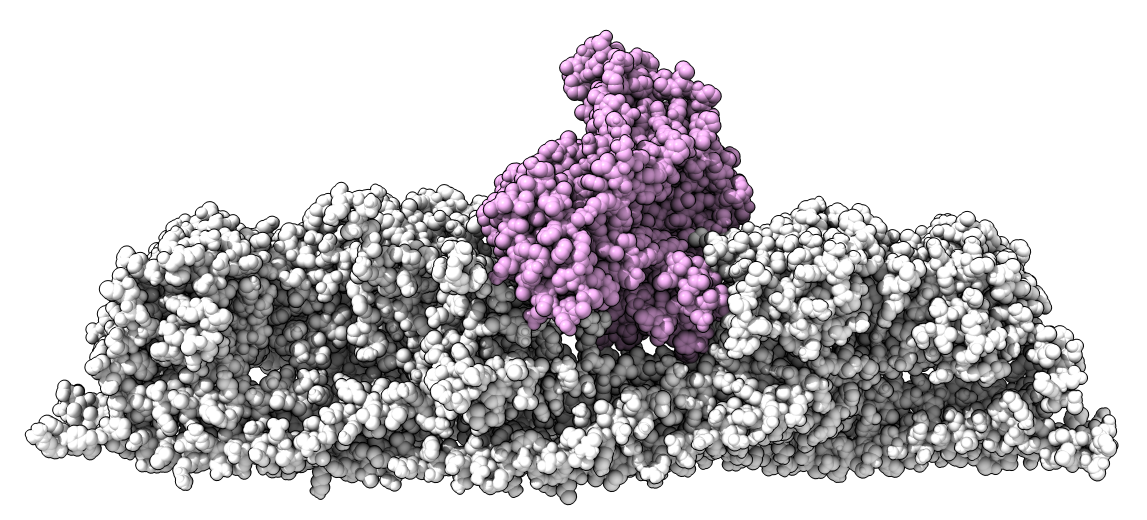

**Cluster II:** Docking resulted in 756 models clustered by Balanced score:

| **Cluster** | **Members** | **Representative** | **Weighted Score** |
| --- | --- | --- | --- |
| **0** | 108 | Center | -629.93 |
|  |  | Lowest Energy | -792.84 |
| **1** | 50 | Center | -621.92 |
|  |  | Lowest Energy | -909.99 |
| **2** | 44 | Center | -610.54 |
|  |  | Lowest Energy | -844.83 |
| **3** | 36 | Center | -622.79 |
|  |  | Lowest Energy | -740.53 |
| **4** | 35 | Center | -621.39 |
|  |  | Lowest Energy | -886.62 |
| **5** | 32 | Center | -625.24 |
|  |  | Lowest Energy | -684.01 |
| **6** | 31 | Center | -1042.04 |
|  |  | Lowest Energy | -1042.04 |
| **7** | 29 | Center | -640.09 |
|  |  | Lowest Energy | -781.66 |
| **8** | 25 | Center | -750.34 |
|  |  | Lowest Energy | -808.43 |
| **9** | 22 | Center | -620.36 |
|  |  | Lowest Energy | -777.11 |
| **10** | 22 | Center | -596.84 |
|  |  | Lowest Energy | -684.66 |
| **11** | 20 | Center | -612.65 |
|  |  | Lowest Energy | -754.48 |
| **12** | 20 | Center | -622.51 |
|  |  | Lowest Energy | -811.62 |
| **13** | 20 | Center | -730.75 |
|  |  | Lowest Energy | -730.75 |
| **14** | 20 | Center | -827.28 |
|  |  | Lowest Energy | -865.78 |
| **15** | 20 | Center | -665.32 |
|  |  | Lowest Energy | -675.52 |
| **16** | 19 | Center | -608.51 |
|  |  | Lowest Energy | -782.17 |
| **17** | 19 | Center | -603.90 |
|  |  | Lowest Energy | -671.36 |
| **18** | 18 | Center | -589.61 |
|  |  | Lowest Energy | -766.37 |
| **19** | 17 | Center | -594.25 |
|  |  | Lowest Energy | -678.66 |
| **20** | 17 | Center | -658.58 |
|  |  | Lowest Energy | -713.23 |
| **21** | 16 | Center | -616.23 |
|  |  | Lowest Energy | -709.67 |
| **22** | 15 | Center | -620.57 |
|  |  | Lowest Energy | -751.75 |
| **23** | 15 | Center | -598.43 |
|  |  | Lowest Energy | -696.27 |
| **24** | 14 | Center | -671.47 |
|  |  | Lowest Energy | -766.25 |
| **25** | 14 | Center | -621.03 |
|  |  | Lowest Energy | -788.16 |
| **26** | 13 | Center | -606.52 |
|  |  | Lowest Energy | -662.43 |
| **27** | 12 | Center | -644.04 |
|  |  | Lowest Energy | -794.93 |
| **28** | 11 | Center | -606.44 |
|  |  | Lowest Energy | -686.00 |
| **29** | 10 | Center | -587.69 |
|  |  | Lowest Energy | -790.54 |

After manual evaluation of five models, the model that satisfies six restraints out of nine was selected.

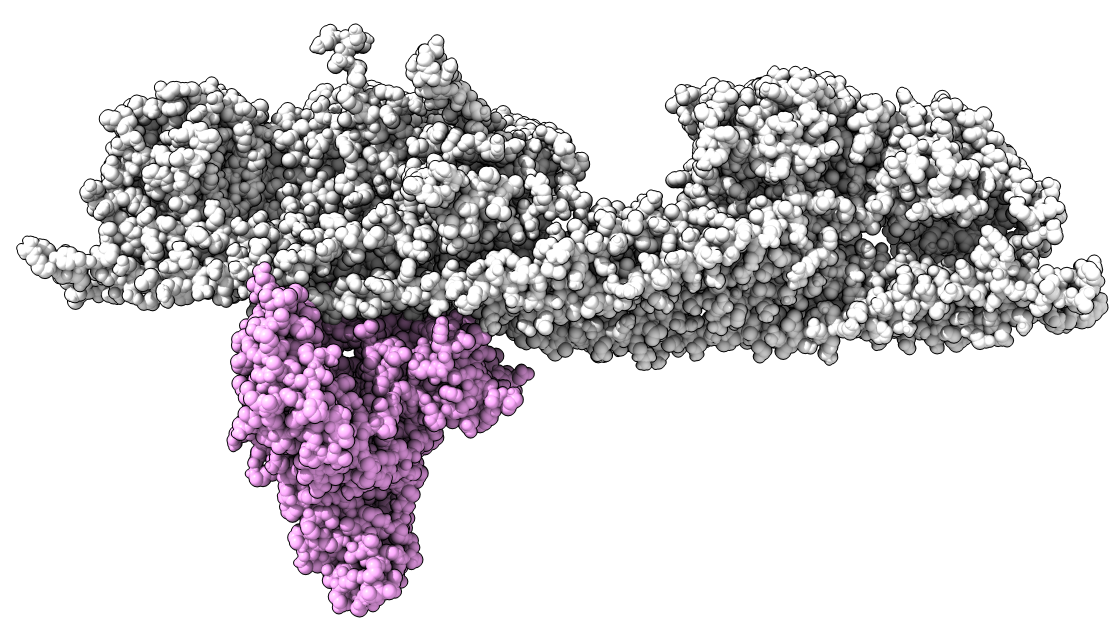

Eventually, 13 out 16 initial restraints were validated. An additional three restraints can be mapped within the defined distance constraint on a model from Cluster I. This model has slightly higher balanced scores than the selected, but satisfies only 9 restraints instead of 11 and was not therefore not selected as the best solution.

**Supplementary Table 5:** Reported plasmin cleavage sites that can be mapped onto the structural model of the fibrin clot.

| Site | Reference | Hindered by HSA | Comment |
| --- | --- | --- | --- |
| αArg’123 | Walker and Nesheim(26) | Yes | - |
| αLys’225 | Henschen et. al.(27) | No | - |
| αLys’249 | Henschen et. al.(27) | No | - |
| αLys’602 | Henschen et. al.(27) | Yes | - |
| βLys’72 | Walker and Nesheim(26) | No | - |
| βLys’163 | Walker and Nesheim(26) | No | - |
| γLys’88 | Walker and Nesheim(26) | Yes | - |
| γLys’111 | Walker and Nesheim(26) | No | Hindered by laterally aggregating coiled-coil. Less preferred than γLys’88. |

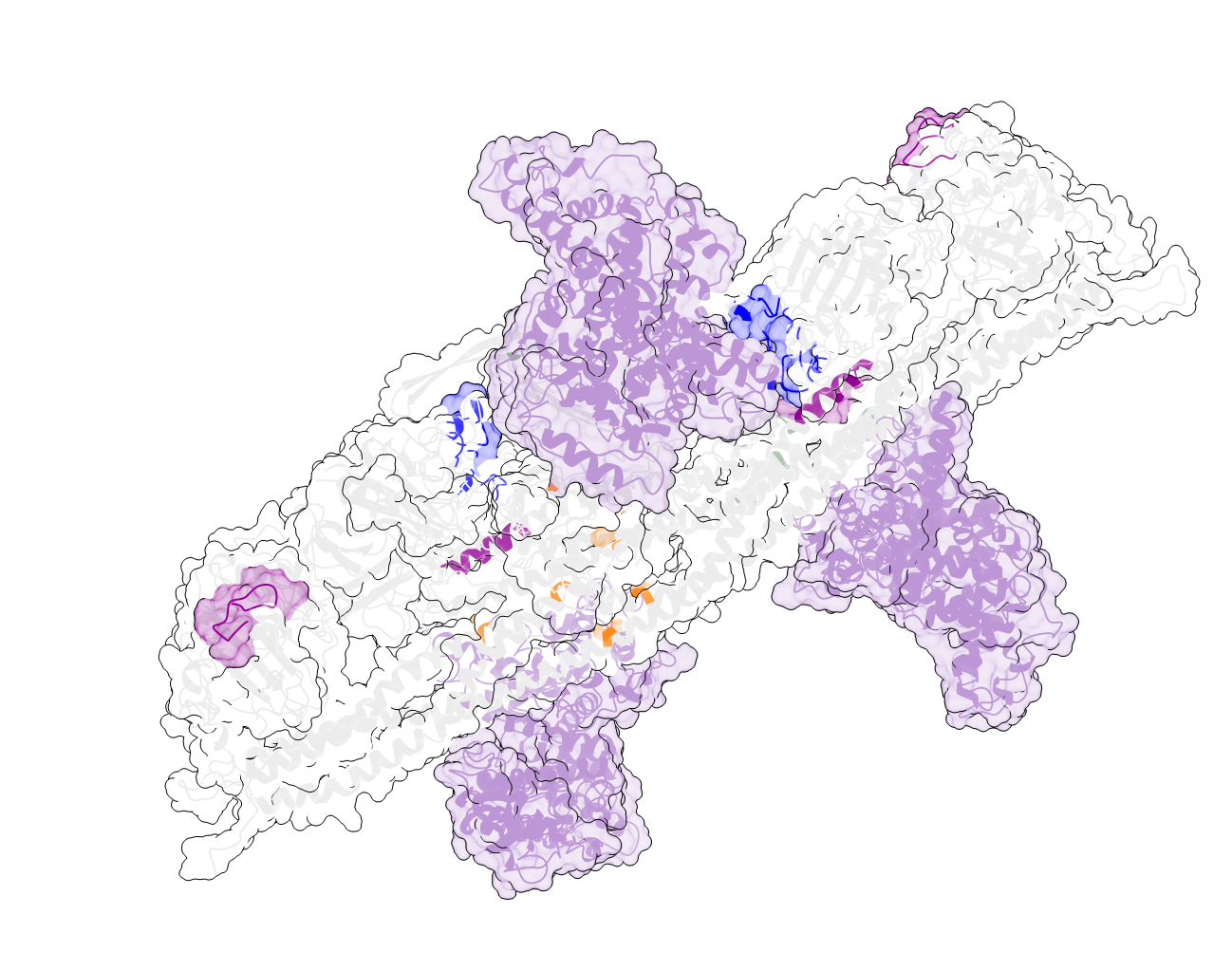

**Supplementary Figure 10:** Suggested placement of three HSA molecules on the fibrin clot. Cluster I places albumin within the cavity on the clots, while Cluster II positions it on the exposed region. As detected crosslinks can be applied to each copy, it is possible to suggest the placement of a third albumin molecule on the exposed region. From our data, we can place 3 copies of albumin per 2 copies of fibrinogen proteins – very close to the stoichiometry suggested from the iBAQ data.
